# Parkin coordinates mitochondrial lipid remodeling to execute mitophagy

**DOI:** 10.1101/2020.02.28.970210

**Authors:** Chao-Chieh Lin, Jin Yan, Meghan D. Kapur, Kristi L. Norris, Cheng-Wei Hsieh, Chun-Hsiang Lai, Nicolas Vitale, Kah-Leong Lim, Ziqiang Guan, Jen-Tsan Chi, Wei-Yuan Yang, Tso-Pang Yao

## Abstract

Mitochondrial failure caused by Parkin mutations contributes to Parkinson’s disease. Parkin binds, ubiquitinates, and targets impaired mitochondria for autophagic destruction. Robust mitophagy involves peri-nuclear concentration of Parkin-tagged mitochondria, followed by dissemination of juxtanuclear mitochondrial aggregates, and efficient sequestration of individualized mitochondria by autophagosomes. Here, we report that the execution of complex mitophagic events requires active mitochondrial lipid remodeling. Parkin recruits phospholipase D2 to the depolarized mitochondria and generate phosphatidic acid (PA). Mitochondrial PA is subsequently converted to diacylglycerol (DAG) by Lipin-1 phosphatase-a process that further requires mitochondrial ubiquitination and ubiquitin-binding autophagic receptors, NDP52 and Optineurin. We show that Optineurin transports, via Golgi-derived vesicles, a PA-binding factor EndoB1 to ubiquitinated mitochondria, thereby facilitating DAG production. Mitochondrial DAG activates both F-actin assembly to drive mitochondrial individualization, and autophagosome biogenesis to efficiently restrict impaired mitochondria. Thus Parkin, autophagic receptors and the Golgi complex orchestrate mitochondrial lipid remodeling to execute robust mitophagy.

## Introduction

Mutations in the ubiquitin E3 ligase, Parkin (PARK2), are the most common cause of early onset Parkinson’s disease (AR-JP) ^1^. Genetic and biochemical evidence have shown that Parkin, working in conjunction with the mitochondrial kinase, PINK1 (PARK6), preserves a healthy mitochondrial population^2–4^. Parkin and PINK1 maintain mitochondrial integrity, at least in part, by recognizing and targeting the impaired and de-energized mitochondria to autophagy-a double membrane vesicles that deliver mitochondria to lysosomes-for destruction ^5, 6. In a basic model, stabilized PINK1 recruits Parkin to depolarized mitochondria, where Parkin catalyzes extensive ubiquitination to initiate mitophagy7–9^. Mitochondrial ubiquitination serves at least two main functions: marking mitochondrial outer membrane proteins for proteasome-mediated degradation^10, 11^; recruiting autophagosomes to sequester and deliver ubiquitin-tagged mitochondria to lysosomes for degradation ^7, 12^. While the exact contribution of the proteasome to mitophagy remains uncertain ^13^, on the autophagy front, mitochondrial ubiquitination has been shown to recruit multiple ubiquitin-binding autophagic receptors, including Optineurin (OPTN), NDP52 and p62/SQSTM1^12, 14, 15^, which bring together ubiquitinated mitochondria and autophagosomes through their binding of LC3-a protein being conjugated to phosphatidylethanolamine in the autophagosome membrane ^16^. Detailed functional analyses, however, have also revealed a more complex picture of this model. Whereas OPTN and NDP52 are required for mitophagy^14, 15^, p62 appears to be dispensable in some studies but not others ^7, 12, 17, 18^. Instead, p62 acts to coalesce ubiquitinated mitochondria into large aggregates at the perinuclear region-a dramatic event yet without a clearly defined function^17, 18^. Recent evidence further indicates that autophagosomes are actually synthesized “de novo” at or around mitochondria tagged for destruction^15, 19, 20^. How Parkin and different autophagic receptors coordinate local autophagosome biogenesis around condemned mitochondria to efficiently execute mitophagy remains to be fully elucidated.

Intriguingly, when subject to massive uncoupling by CCCP, the large mitochondrial populations tagged by Parkin are recognized and actively concentrated by microtubule-based motors and form large perinuclear aggregates or inclusions ^12, 21^. Perinuclear mitochondrial inclusions have also been observed in neurons from PD patients ^22, 23^, indicating that this process is physiologically relevant. Juxtanuclear mitochondrial aggregates, or termed mito-aggresomes, are subsequently dispersed into smaller units prior to their sequestration by autophagosomes (mitophagosomes) and eventual degradation^11^. Electron microscopy has revealed that some perinuclear mitochondria concentrated by Parkin and PINK1 are fused ^21^. These findings imply that dissemination of perinuclear mitochondrial aggregates would require active “de-aggregation” and “fission” of mitochondria. Thus, the disposal of impaired mitochondria involves elaborate temporal and spatial regulation orchestrated by Parkin. How Parkin coordinates these events to achieve robust mitophagy is not well understood.

In this report, we present evidence that the recruitment of Parkin to depolarized mitochondria initiates a series of lipid remodeling events, characterized by the sequential production of mitochondrial phosphatidic acid (PA) and diacylglycerol (DAG). Using fluorescent PA and DAG binding reporters, we present evidence that Parkin recruits a phospholipase, PLD2, to the depolarized mitochondria to generate PA, while activates the PA phosphatase Lipin-1-dependent DAG production through mitochondrial ubiquitination that recruits ubiquitin-binding autophagic receptors, OPTN and NDP52. We present evidence that OPTN and NDP52, via Golgi-derived vesicles, deliver a lipid-remodeling factor EndoB1 to the ubiquitinated mitochondria and stimulate DAG synthesis. Functionally, mitochondrial DAG drives both the individualization of juxtanuclear mitochondrial aggregates by stimulating F-actin assembly on mitochondria, and the production of autophagosomes, thereby coupling the preparation of autophagible mitochondria units with autophagosome sequestration. Our findings uncover local mitochondrial lipid remodeling as a novel and critical mechanism that ensures robust mitophagy, and identify the Golgi complex as an important component of mitochondrial QC machinery.

## Results

### Parkin stimulates mitochondrial PA and DAG accumulation

PINK1 and Parkin have been shown to promote mitochondrial fusion in response to moderate mitochondrial stresses induced by low-dose CCCP ^24^. In addition to mitofusin (MFN)- and Drp1-related GTPases, mitochondrial dynamics is affected by membrane lipid composition, in which mitochondrial phosphatidic acid (PA) and diacylglycerol (DAG) stimulate fusion and fission respectively ^25, 26^. We investigated whether mitochondrial PA or DAG is involved in Parkin-mediated mitochondrial fusion. To this end, we employed specific fluorescent reporters that bind PA (EGFP-Raf1-PABD, referred as PA reporter ^27^) or DAG (C1bδ-CFP/YFP, referred as YFP-DAGR) to monitor PA and DAG production ^26, 28^. However, when Parkin-expressing cells were challenged by low-dose CCCP (1 μM), no discernible change was observed for either the PA reporter or DAGR (not shown). Unexpectedly, when these cells were challenged by high-dose CCCP (10 μM) to activate global mitophagy, prominent accumulation of the PA-reporter on Parkin-tagged mitochondria were observed, as early as 1 hour after CCCP treatment (Fig. 1a, quantified in Fig. 1c). A different PA-binding reporter (Spo20p-GFP) ^29^ showed similar mitochondrial translocation in response to high-dose CCCP treatment (Supplementary Fig. 1a). Mitophagy activated by antimycin/oligomycin treatment similarly induced concentration of the PA reporter on mitochondria. (Supplementary Fig. 1b). Using mitochondria-targeted photosensitizer KillerRed to damage a defined population of mitochondria^20^, the PA reporter was again detected on Parkin-positive mitochondria (Supplementary Fig. 1d). Collectively, these findings indicate that PA accumulates on impaired mitochondria tagged by Parkin.

**Fig. 1.**
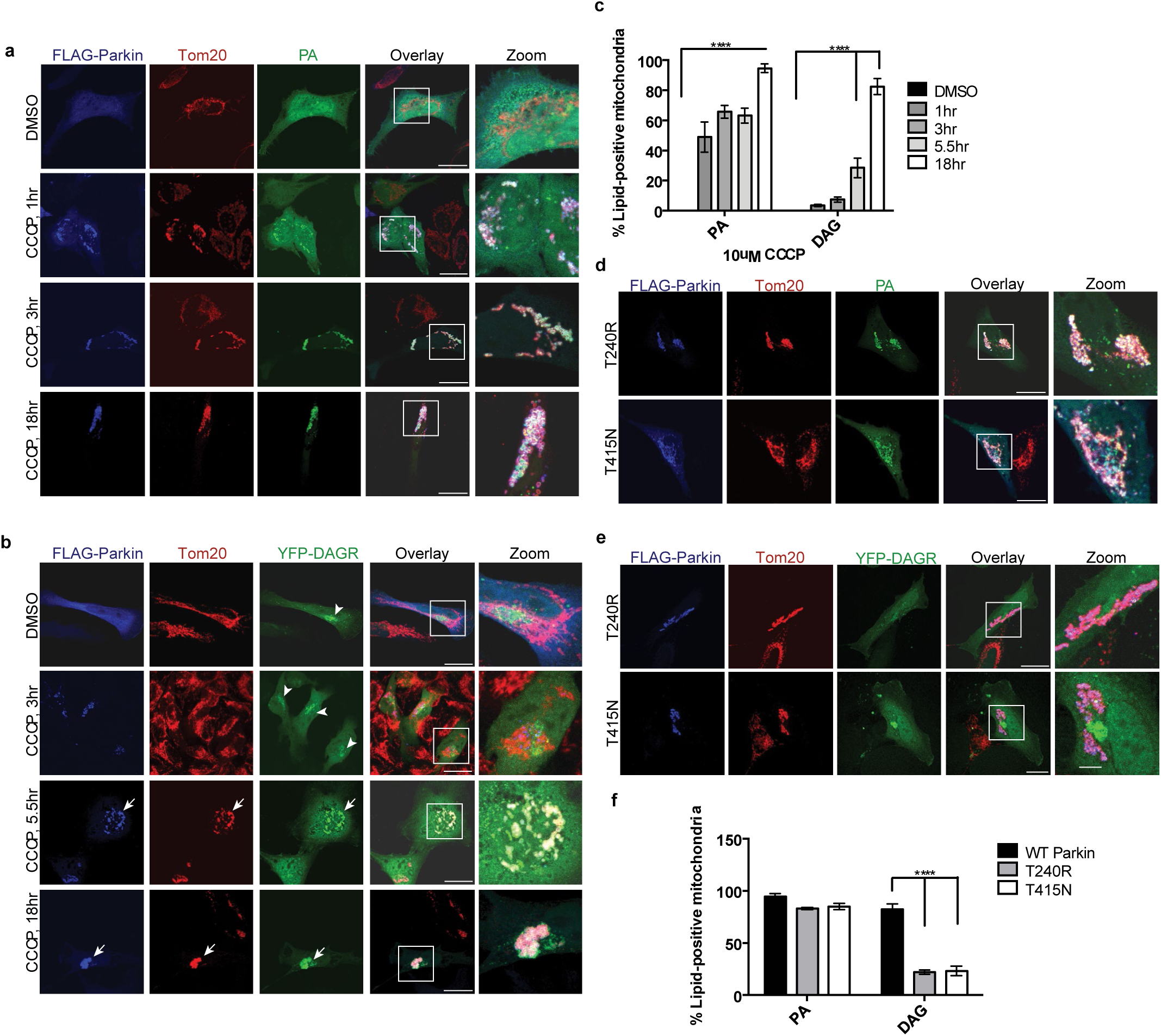
Parkin stimulates PA and DAG accumulation on depolarized mitochondria. HeLa cells were transfected with a WT or mutant Parkin plasmid (FLAG), shown in blue, along with reporters for either PA or DAG (see Materials and Methods), shown in green. DMSO or 10 µM CCCP was added for the indicated periods of time. Mitochondria were visualized via anti-Tom20 staining (red). **a** PA accumulated on Parkin-positive mitochondria after 1 hour of 10uM CCCP treatment, with a more dramatic increase by 3hr and 18 hr. **b** DAG (diacylglycerol) accumulated on Parkin-positive mitochondria after 5.5 hr of 10uM CCCP treatment. **c** Quantification of imaging experiments shown in A and B. **d** E3 ligase-deficient Parkin mutants (T240R and T415N) have no effect on PA accumulation (green) on depolarized, Parkin-positive mitochondria. **e** DAG accumulation depends upon intact Parkin E3 ligase activity. Scale bar = 25µm and zoom is 3x. **f** Quantification of imaging experiments shown in D and E. Bars represent mean with SEM from three independent experiments; two-way ANOVA analysis was performed for statistical analysis (***P<0.0001).

To assess whether PA on Parkin-tagged mitochondria is further processed to DAG, we employed a high-affinity DAG reporter (YFP-DAGR). Under basal conditions YFP-DAGR mainly labeled perinuclear structures corresponding to the Golgi apparatus, which contains abundant DAG ^30^ (Fig. 1b, arrowhead). Upon mitophagy activation by either CCCP or antimycin/oligomycin treatment, dramatic recruitment of YFP-DAGR to Parkin-tagged mitochondria was observed (Fig. 1b-1c and Supplementary Fig. 1c). Neither YFP-DAGR nor the PA reporter accumulated on mitochondria in the absence of Parkin (not shown); however, both reporters were detected on mitochondria targeted for mitophagy in SH-SY5Y cells, which express endogenous Parkin (Supplementary Fig. 1e-f). Altogether, the lipid reporter assays indicate that Parkin stimulates PA and DAG production on mitochondria tagged for autophagy.

### Mitochondrial DAG production requires Parkin E3 ligase activity

Parkin activates mitophagy by catalyzing mitochondrial ubiquitination. We assessed two AR-JP associated Parkin mutants, T240R and T415N, which are deficient in the ubiquitin E3 ligase activity and cannot support mitophagy ^12^. As shown in Fig. 1d, neither mutations affected the mitochondrial translocation of the PA-reporter. In contrast, mitochondrial YFP-DAGR accumulation was strongly inhibited (Fig. 1e, quantified in 1f). Thus, mitochondrial ubiquitination is required for mitochondrial DAG, but not PA, production. These findings suggest that mitochondrial PA production is initiated by the recruitment of Parkin while DAG production further requires Parkin E3 ligase activity.

### PLD2 and Lipin-1 catalyze sequential production of mitochondrial PA and DAG

We next determined how Parkin orchestrates mitochondrial lipid remodeling. Phospholipase D is the primary enzyme that produces PA. A mitochondrial resident phospholipase D, Mito-PLD (PLD6), generates mitochondrial PA by hydrolyzing cardiolipin ^25^. However, PA reporter accumulated normally on Parkin-tagged mitochondria in Mito-PLD knockout fibroblasts in response to CCCP treatment (data not shown). We next investigated whether Parkin recruits a cytosolic PLD to depolarized mitochondria and catalyze PA production. Of two widely expressed PLD members, we found that PLD2 (Fig. 2a), but not PLD1 (Supplementary Fig. 2a), translocated onto Parkin-tagged mitochondria upon treatment with CCCP or oligomycin/antimycin (Supplementary Fig. 2b). Immunoblotting confirmed that endogenous PLD2, along with Parkin, became enriched in the mitochondrial fraction in response to CCCP treatment (Fig. 2b). To determine the role of PLD2 on mitochondrial PA production, we inhibited PLD2 by a specific inhibitor, VU 0364739^31^, or by siRNA-mediated knockdown. Importantly, mitochondrial accumulation of the PA reporter was markedly suppressed by either pharmacological inhibition (Fig. 2c-d) or siRNA-mediated knockdown of PLD2 (Supplementary Fig. 2d-e). PLD2 inhibition also markedly prevented mitochondrial YFP-DAGR accumulation (Supplementary Fig. 2f). These findings indicate that Parkin stimulates mitochondrial PA production by recruiting PLD2, and that PLD2-generated PA is a precursor to mitochondrial DAG.

**Fig. 2.**
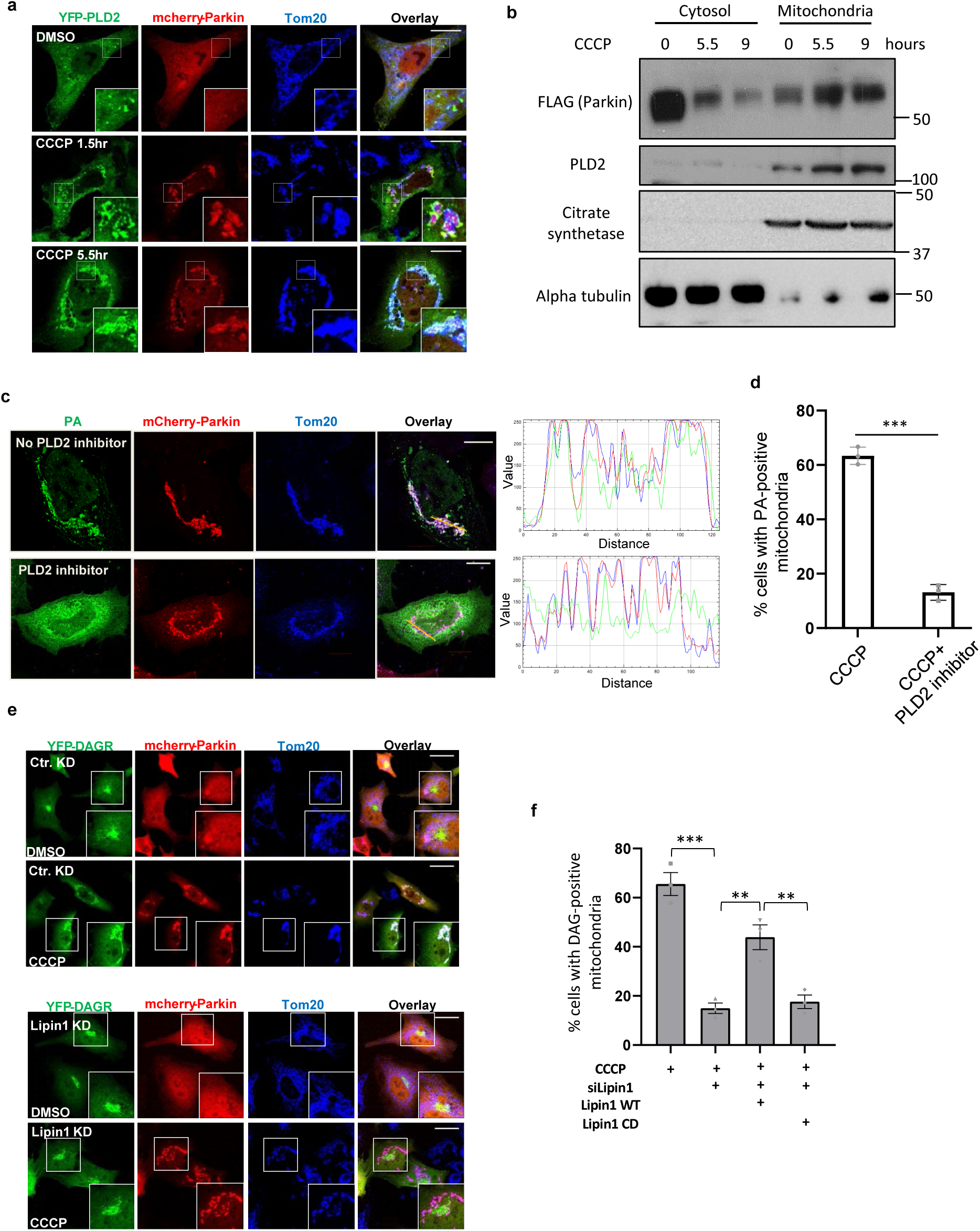
PLD2 and lipin1 work in tandem to stimulate mitochondrial PA and DAG production. **a** HeLa cells were transfected with expression plasmids for Parkin (mCherry-Parkin, Red) and YFP-PLD2 (Green) and treated with DMSO or 10 µM CCCP for the indicated time. Mitochondria were visualized via anti-Tom20 staining (blue). Note that YFP-PLD2 accumulated on Parkin-positive mitochondria after CCCP treatment. Scale bar = 25µm. **b** HEK-293T cells were transfected with Parkin-FLAG and DAGR-YFP followed by CCCP treatment for indicated times. Mitochondrial and cytosolic fractions were purified and analyzed by Western blots by indicated antibodies. Citrate synthetase and α-tubulin were used as a mitochondrial and cytosolic marker, respectively. Note that PLD2 and Parkin levels were elevated in the mitochondrial fraction in response to CCCP treatment. **c** Hela cells were co-transfected with mcherry-Parkin and the PA reporter, followed by CCCP treatment alone or with a PLD2 inhibitor VU 0364739 (3 µM) for 5.5 hrs. Line scan analysis (Image J software) corresponding to the line drawn in the images indicate colocalization between the PA reporter (green) and mitochondria (red). Note that VU 0364739 suppressed mitochondrial PA accumulation. Scale bar = 10 µm. **d** Quantification of the numbers of cells with the PA reporter positive mitochondria shown in **c**. Asterisks indicate statistical significance (***P<0.001, Student’s t-test). **e** Hela cells were transfected with a lipin1 siRNA, followed by the DAG reporter and mCherry-Parkin, treated with CCCP (10 µM) for 9 hrs and then subject to image analysis. Scale bar = 25 um and zoom is 4x. **f** The numbers of cells with DAG-reporter positive mitochondria shown in **e** and in Lipin-1 knockdown cells transfected with siRNA-resistant wildtype (WT) and catalytically dead (CD; D712E;D714E) mutant cDNA. Note that Lipin1 KD reduced mitochondrial DAG accumulation, which can be significantly restored by the re-expression of WT, but not CD mutant, Lipin-1. The graph shows the means with SEM (error bars) from three independent experiments. Asterisks indicate statistical significance by one-way ANOVA (**P< 0.01, ***P<0.001). Source data are provided as a Source Data file.

To determine how PA is converted to DAG, we knocked down Lipin-1, a major cytosolic phosphatase that dephosphorylates PA to DAG ^26, 32^. Lipin-1 knockdown led to a dramatic loss of DAGR-YFP on Parkin-tagged mitochondria (Fig. 2e, Lipin-1 KD) while the PA reporter was not affected (Supplementary Fig. 2h). Re-expression of a siRNA-resistant WT Lipin-1 in Lipin-1 KD cells significantly restored mitochondrial DAGR-YFP while a catalytic inactive mutant (Lipin-1 CD) was ineffective (Fig. 2f). Together, these findings demonstrate that PLD2 and Lipin-1 work in tandem to generate PA and DAG on mitochondria tagged by Parkin.

### DAG coordinates F-actin-dependent mitochondrial dissemination and mitophagy

Compared to WT control, we found that Lipin-1 knockdown cells were less efficient in mitochondrial clearance, as shown by the significant retention of mitochondria following CCCP treatment (Fig. 3a, bottom panels, quantified in Fig. 3b). Immunoblotting analysis confirmed that autophagy-dependent degradation of multiple mitochondrial markers was impaired in lipin-1 KD cells, whereas proteasome-mediated degradation of MFN1 was unaffected (Fig. 3c and Supplementary Fig. 2i). PLD2 inhibition, which suppressed the production of mitochondrial PA, the precursor to DAG, also reduced the efficiency of mitophagy (Supplementary Fig. 2j). These results support the conclusion that mitochondrial DAG production is important for Parkin-mediated mitophagy.

**Fig. 3.**
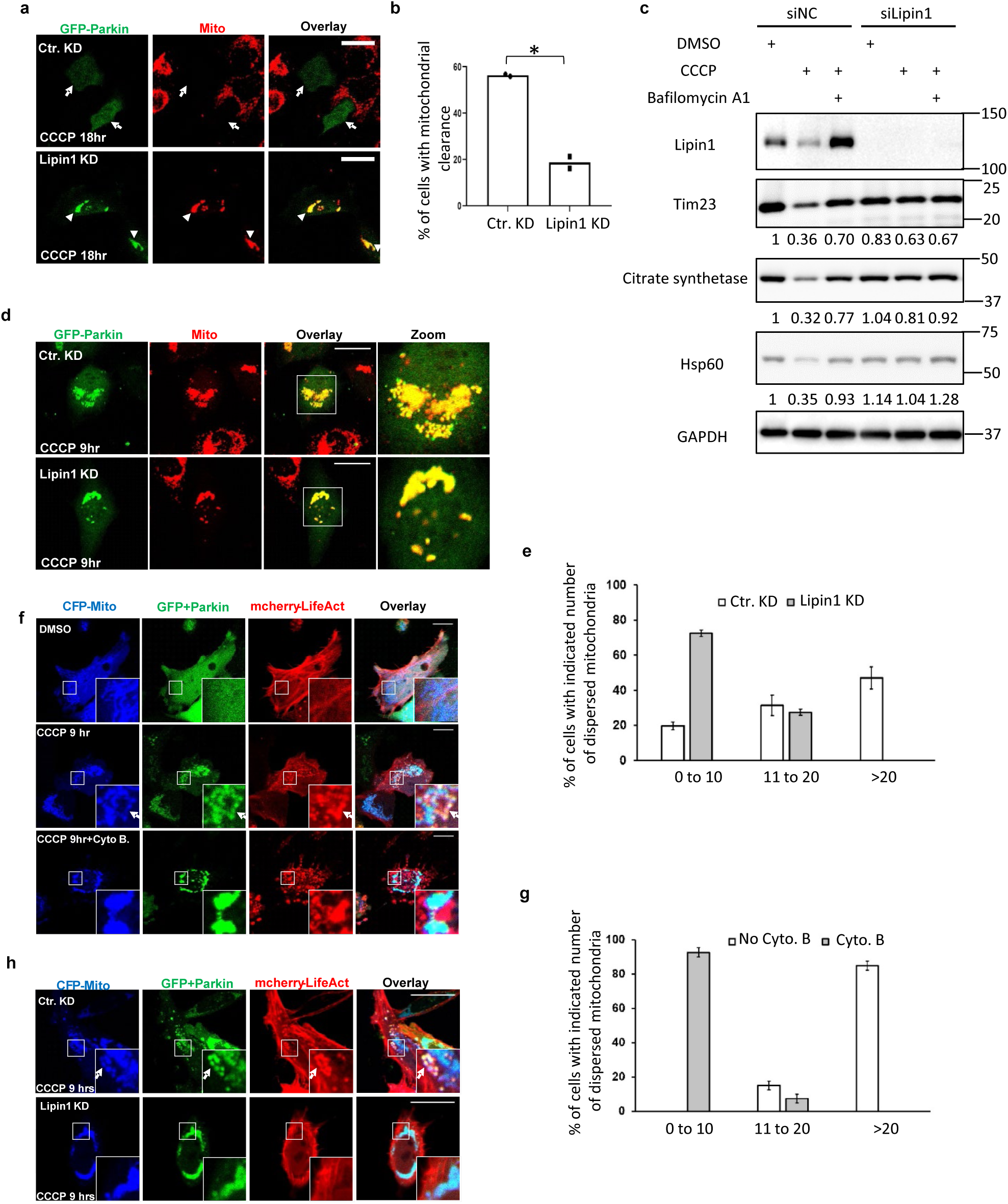
Lipin1 regulates mitophagy and F-actin-dependent mitochondrial dispersion. **a,b** Hela cells were transfected with a Lipin1 or control (cKD) siRNA followed by a GFP-Parkin expression plasmid. Transfected cells were treated with CCCP at 10 µM for 18 hrs, and subject to immuno-staining. **b** Quantification of Parkin-positive cells in **a** that have lost a majority of mitochondria (marked by TOM20, arrows) from two independent experiments. Asterisks indicate statistical significance (*P<0.05, Student’s t-test). Note that mitochondrial clearance is reduced in Lipin-1 knockdown cells (arrowheads in **a**). **c** Control or lipin1 knockdown Hela cells stably expressing parkin-mCherry were treated with DMSO or CCCP (10 µM for 18 hrs) and Bafilomycin A1 (1μM, lysosomal inhibitor) treatment, followed by immunoblotting with indicated antibodies. Note that lipin1 silencing rescued mitochondrial protein degradation. The band intensity of mitochondrial proteins relative to control untreated conditions were determined by Image J. **d** HeLa cells were treated and processed as described in **a** but with 9 hr CCCP treatment. Small and dispersed mitochondria were prominent in control knockdown cells while this dispersion was suppressed in Lipin-1 knockdown cells. **e** Quantification of the number of dispersed mitochondria, as shown in **d**. **f-h** Control or Lipin-1 knockdown Hela cells were co-transfected with GFP-Parkin, a mitochondrial marker (CFP-Mito), and F-actin marker (RFP-LifeAct) followed by CCCP treatment alone or with Cytochalasin B at 10 µM, as indicated. Live cell images were then acquired and analyzed. Note the marked co-localization of RFP-LifeAct and Parkin-positive mitochondria in control but not Cytochalasin B treated **f** or Lipin-1 KD cells **h**. **g** Quantification of the effect of Cytochalasin B on mitochondrial dispersion from **f**. Scale bar = 25 µM and zoom is 3x **d** or 12x **f** or 7x **h**. Source data are provided as a Source Data file.

During mitophagy, individual mitochondria became dispersed from the perinuclear aggregates prior to their sequestration by autophagosomes ^11^. Indeed, we observed many small Parkin-positive mitochondria separated from the perinuclear cluster during mitophagy (Fig. 3d, Top Panels). Strikingly, in Lipin-1 KD cells, significantly fewer “individualized and dispersed” mitochondria were observed. Instead, mitochondria frequently remained as large perinuclear aggregates (Fig. 3d, Bottom Panels; quantified in Fig. 3e). These findings indicate that Lipin-1 is required for the dispersion of mitochondrial aggregates prior to autophagic sequestration.

Autophagic clearance of perinuclear protein inclusions, the aggresomes, requires active de-aggregation implemented by the actin cytoskeleton^33^. Because DAG is a known activator of F-actin assembly on the plasma membranes and endosomes^34, 35^, we asked whether Lipin-1-dependent DAG production activates mitochondrial de-aggregation by promoting F-actin assembly. To this end, we first used an F-actin binding probe, RFP-LifeAct^36^, to assess if F-actin was assembled around mitochondrial aggregates. As shown in Fig. 3f, prominent LifeAct-positive signals were detected on de-aggregated mitochondria (Middle Panels), suggesting that F-actin is involved in mitochondrial de-aggregation. Supporting this hypothesis, inhibition of F-actin assembly by cytochalasin B (Cyto B) markedly suppressed the appearance of RFP-LifeAct positive mitochondria (Fig. 3f, Bottom Panels) and the dispersion of mitochondrial aggregates (Quantified in Fig. 3 g). Importantly, Lipin-1 knockdown also suppressed the appearance of LifeAct positive-mitochondria (Fig. 3h). These results support the conclusion that Lipin-1-dependent production of mitochondrial DAG stimulates local F-actin assembly to divide and disperse perinuclear mitochondrial aggregates for autophagic degradation.

### Lipin1-dependent DAG stimulates autophagosome production

We noticed that abundant accumulation of YFP-DAGR on mitochondria apparently retarded mitochondrial clearance, as indicated by the continuous accumulation of YFP-DAGR positive mitochondria after prolonged CCCP treatment (Fig. 1c, 18 hr). This finding is consistent with the notion that high affinity DAGR can mask mitochondrial DAG from the downstream effectors. To follow the fate of DAG at a later stage of mitophagy, we employed another DAG reporter with a lower binding affinity, C1(2)-mRFP (referred as RFP-DAGR^37^). Under basal condition, RFP-DAGR also mainly labeled the Golgi complex (Fig. 4a, top panels, arrowhead) positive for a trans-Golgi marker TGN46 (Supplementary Fig. 3a). Interestingly, upon CCCP treatment, many RFP-DAGR positive vesicles encompassing dispersed parkin-tagged mitochondria were observed (Fig. 4b, Top Panels). Similar structures were also detected using the YFP-DAGR reporter expressed at a lower concentration (Supplementary Fig. 3b). Importantly, RFP-DAGR - labeled vesicles are frequently positive for the autophagosome marker LC3 (Fig. 4b) suggesting that mitochondrial DAG is also involved in the production of autophagosomes that restrict damaged mitochondria (i.e. mitophagosomes). Supporting this possibility, the abundance of LC3-positive vesicles was markedly reduced in Lipin-1 KD cells after CCCP treatment (Fig. 4b, bottom panels, and quantified in Fig. 4c). The LC3 signals remained in Lipin-1 KD cells were typically small and did not contain mitochondria (Fig. 4b, Zoomed Insets). As expected, mitophagy-associated RFP-DAGR induction was suppressed in Lipin-1 KD (Fig. 4b, bottom panels, Red). Collectively, these findings indicate that Lipin-1 and mitochondrial DAG production stimulate mitophagosome production.

**Fig. 4.**
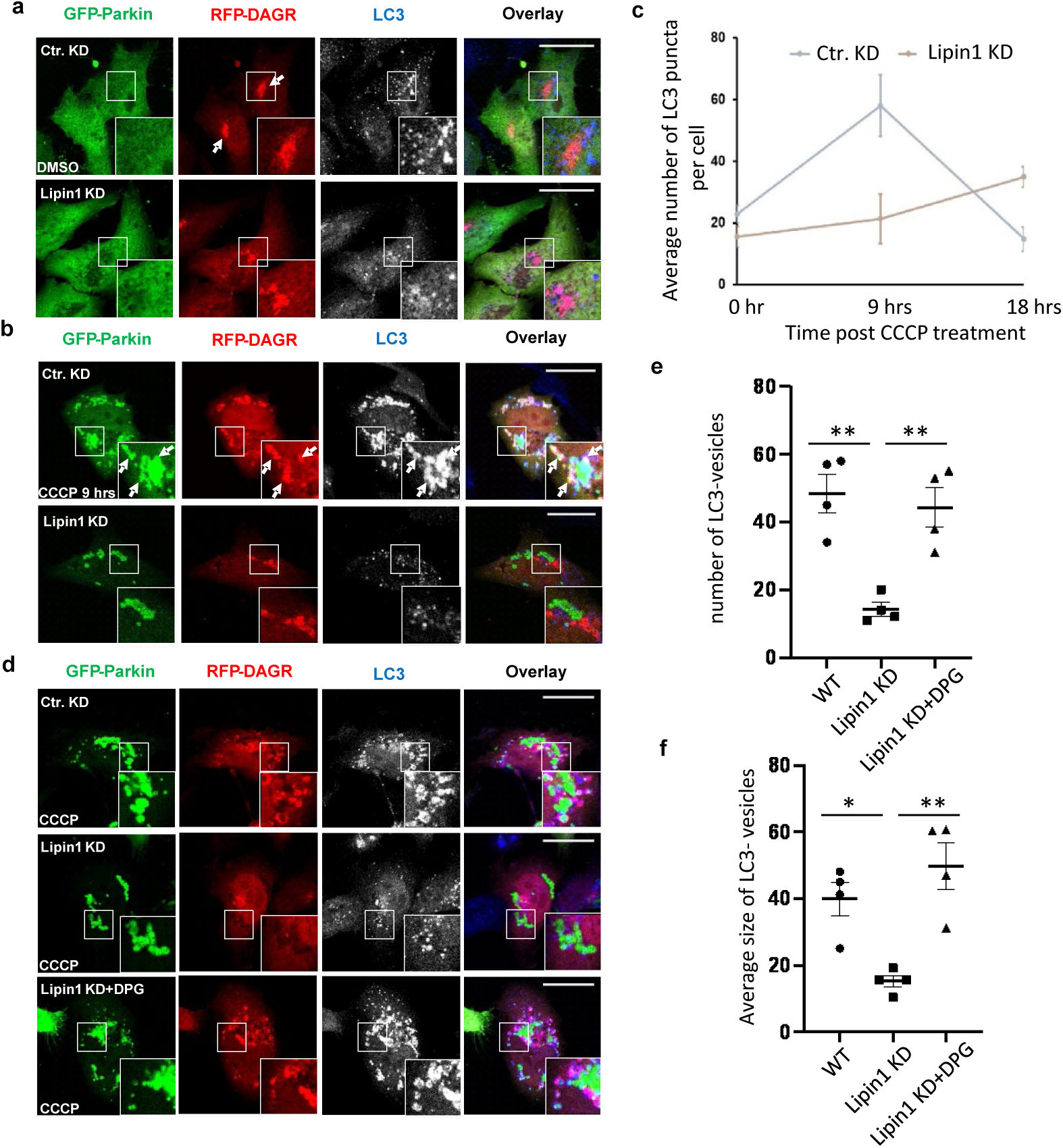
Lipin-1-mediated mitochondrial DAG production is required for mitophagosome production. Hela cells were transfected with control or a Lipin1 siRNA followed by the expression plasmids of GFP-Parkin and a DAG reporter (RFP-DAGR). **a,b** Transfected cells were treated with DMSO **a** or CCCP **b** at 10 µM for 9 hr as indicated. Autophagosome formation was assessed by a LC3 antibody. LC3 is pseudo colored in white in single channel and blue in the overlay. **c** Quantification of autophagosome (LC3) from experiment in **a**, where at least 50 cells in three independent experiments were analyzed. **d** Lipin1 knockdown cells were further incubated with 1,2-Dipalmitoyl-sn-glycerol (DPG) at 100 µM in addition to CCCP and assessed for autophagosome formation. Note that CCCP induced DAG-positive autophagosomes that sequester dispersed mitochondria in control knockdown cells and this induction is suppressed in Lipin-1 knockdown cells (9 hr post CCCP). 1,2-Dipalmitoyl-sn-glycerol treatment induced DAG-positive autophagosomes in Lipin-1 knockdown cells. Scale bar = 25 µM and zoom is 5x. **e,f** The number **e** and average size (**f**, arbitrary unit) of LC3-vesicles from indicated samples in **d** were quantified by Image J software. Note that both the number and size of LC3-vesicles in Lipin-1 knockdown cells were much smaller than those in WT cells and these defects were corrected by DPG treatment. Asterisks indicate statistical significance by one-way ANOVA (*P< 0.05, **P<0.01) from four independent experiments. Source data are provided as a Source Data file.

To confirm the role of DAG in mitophagosome production, we supplied Lipin-1 KD cells with a cell permeable 1,2-dipalmitoyl-sn-glycerol (DPG), an effector DAG species produced by Lipin-1^38^. DPG supplement (+DPG Panels) markedly stimulated both the appearance of dispersed mitochondria and large LC3-positive vesicles, many of those contained mitochondria (Fig. 4d, Zoomed Inset; Quantified in Fig. 4e-f). These results indicate that mitochondrial DAG produced by Lipin-1 promotes both mitochondrial dissemination and the production of autophagosomes that sequester dispersed mitochondria.

### Mitochondrial DAG production requires ubiquitin-binding NDP52 and Optineurin

Parkin-dependent mitochondrial DAG production requires its ubiquitin E3 ligase activity (Fig. 1e). Mitochondrial ubiquitination was shown to recruit ubiquitin-binding autophagic receptors OPTN and NDP52^14, 15^. We, therefore, determined if OPTN and NDP52 are involved in mitochondrial DAG production. Consistent with their redundant functions in mitophagy ^15^, individual knockdown of OPTN or NDP52 had no discernable effect on mitochondrial YFP-DAGR accumulation (Supplemental Fig. 4a). Simultaneous knockdown of these autophagic receptors, however, abrogated mitochondrial YFP-DAGR accumulation (Fig. 5a-b). Re-expression of an siRNA-resistant WT OPTN, but not a ubiquitin-binding deficient (E478G) mutant OPTN ^14^, restored mitochondrial DAGR recruitment, supporting a specific and ubiquitin-dependent activity of OPTN in mitochondrial lipid remodeling (Fig. 5b). Consistent with their role in mitochondrial DAG production, OPTN/NDP52 KD also suppressed the formation of RFP-DAGR-positive LC3 vesicles (mitophagosomes, Supplementary Fig. 4b). Importantly, exogenous DPG treatment markedly increased mitophagy in OPTN/NDP52 KD cells (Fig. 5c-d). These findings show that a key function of OPTN and NDP52 in mitophagy is to promote mitochondrial DAG production.

**Fig. 5.**
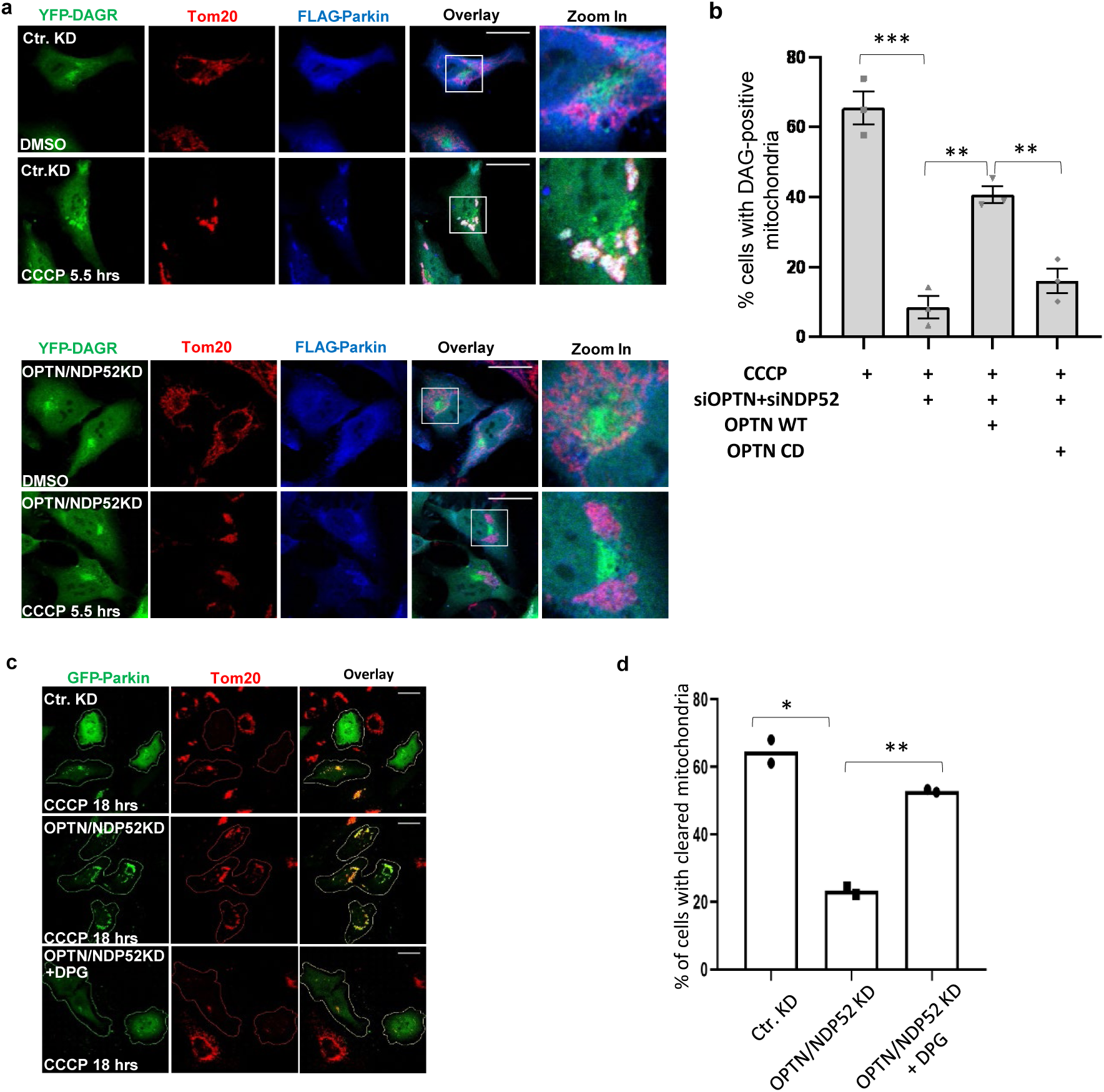
Requirement of OPTN, NDP52 for DAG production and exogenous DAG restores mitophagy. **a** Hela cells were transfected with control or optineurin and NDP52 siRNAs followed by expression plasmids of FLAG-Parkin **a** with the YFP-DAGR. These cells were treated with DMSO or 10 µM CCCP for 5.5 hrs, as indicated. Note that OPTN/NDP52 double knockdown prevented mitochondrial YFP-DAGR accumulation. **b** Hela OPTN/NDP52 knockdown cells were transfected with a siRNA-resistant wildtype or ubiquitin-binding deficient E478G OPTN mutant, as indicated. The percentage of cells with mitochondrial YFP-DAGR following CCCP treatment was quantified. Asterisks indicate statistical significance by one-way ANOVA (**P< 0.01, ***P<0.001) from three independent experiments. **c** Hela OPTN/NDP52 knockdown cells were generated as **a** and incubated with 1,2-Dipalmitoyl-sn-glycerol followed by CCCP treatment for 18 hrs. Note that mitophagy was markedly restored by 1,2-Dipalmitoyl-sn-glycerol in OPTN/NDP52 knockdown cells. **d** Quantification of percentage of cells with cleared mitochondrial in **c**. Asterisks indicate statistical significance (*P<0.05, **P<0.01, Student’s t-test) from two independent experiments. Scale bar = 25 µM and zoom is 3x. Source data are provided as a Source Data file.

### Mitochondrial DAG production requires Golgi associated EndoB1

Although commonly known as autophagic receptors, OPTN and NDP52 were initially characterized as trans-Golgi associated proteins that regulate Golgi vesicle transport^39, 40^. Indeed, fluorescence-tagged OPTN (OPTN-Cherry) appeared as perinuclear vesicle-like structures that were partially co-localized with a trans-Golgi marker, TGN38 (TGN38-GFP) (Fig. 6a, Top Panels). Importantly, TGN38, along with OPTN, became accumulated on Parkin-tagged mitochondria upon CCCP treatment (Fig. 6a, Bottom Panels). These observations suggest that OPTN-TGN38-associated vesicles translocate onto ubiquitinated mitochondria tagged by Parkin. These findings raise the possibility that OPTN might deliver factor(s) from the Golgi complex or associated vesicles to ubiquitinated mitochondria, and thereby stimulate Lipin1-dependent DAG production. We searched for potential Golgi factor(s) that is known to modify membrane lipid composition and regulate mitophagy. One such a candidate identified is Endophilin B1 (EndoB1, also known as Bif-1), which was reported to possess an intrinsic PA-binding activity^41^ and is require for mitophagy^41, 42^. We found that EndoB1 was concentrated on the OPTN-positive vesicles under basal conditions. In response to CCCP, EndoB1 became associated with Parkin- and OPTN-tagged mitochondria (Fig. 6b). Mitochondrial recruitment of EndoB1, along with OPTN, was confirmed by immunoblotting (Fig. 6c). Supporting EndoB1 as a potential cargo delivered to mitochondria by OPTN/NDP52, mitochondrial recruitment of EndoB1 was markedly reduced in OPTN/NDP52 KD cells (Fig. 6d-e). Importantly, knockdown of EndoB1 suppressed the accumulation of YFP-DAGR on Parkin-tagged mitochondria (Fig. 6f-g) but had no effect on PA reporter translocation (Supplementary Fig. 5a). Consistent with its effect on mitochondrial DAG production, EndoB1 knockdown also reduced the production of DAGR-positive LC3 vesicles (Supplementary Fig. 5b), These findings indicate that OPTN and NDP52 promote mitochondrial DAG production and mitophagy, at least in part, by delivering EndoB1 to ubiquitinated mitochondria.

**Fig. 6.**
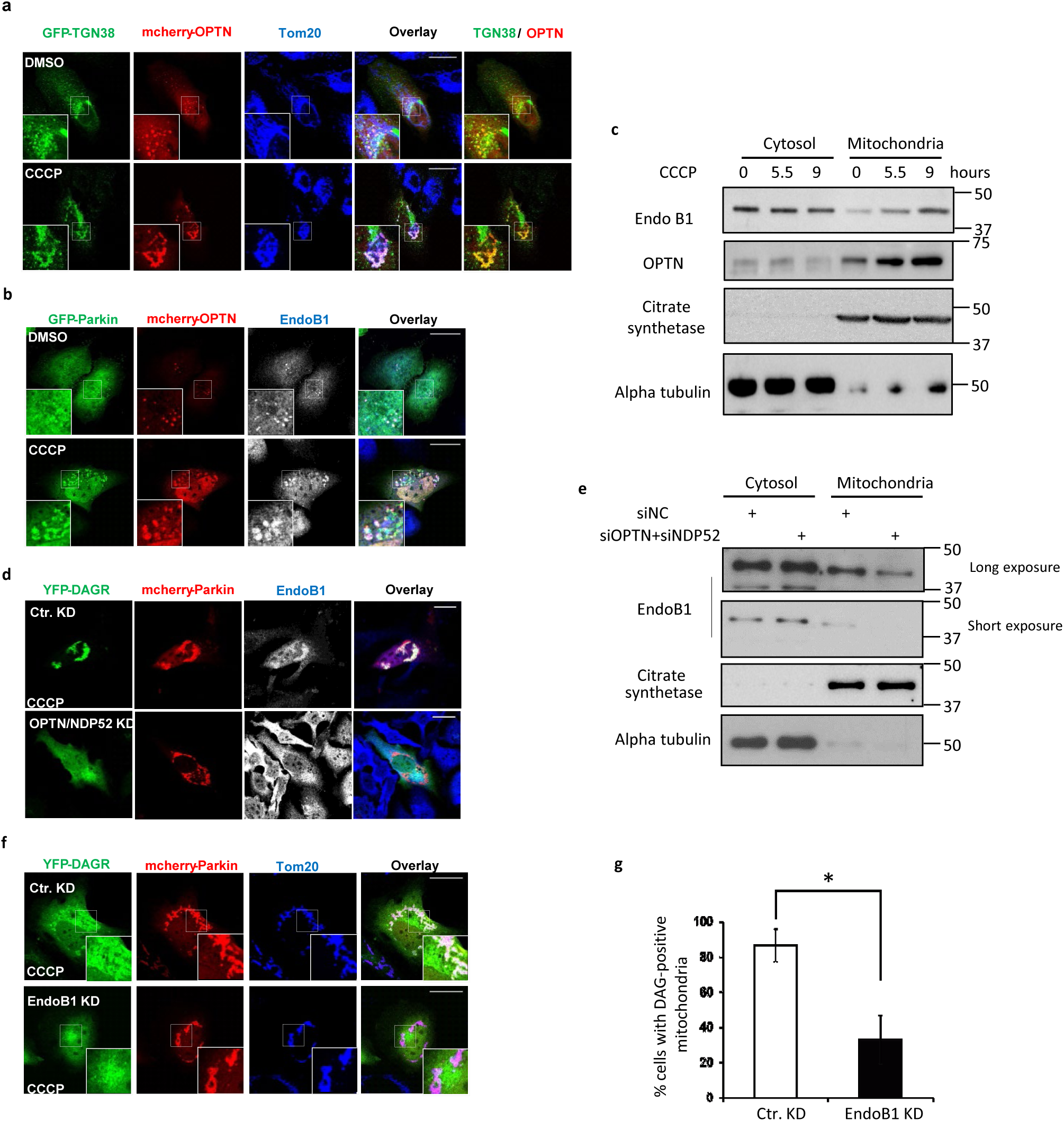
OPTN and NDP52 deliver EndoB1-positive golgi vesicles to ubiquitinated mitochondria for mitochondrial DAG production. **a,b** Hela cells were co-transfected with expression plasmids for GFP-TGN38, mCherry-OPTN and FLAG-Parkin **a** or mCherry-OPTN and GFP-Parkin **b** and treated with CCCP for 5.5 hrs. Mitochondria were visualized by immunostaining with a Tom20 antibody and EndoB1 localization was assessed by an EndoB1 antibody. Note that GFP-TGN38 (**a** bottom panel) and EndoB1 (**b** bottom panel), pseudo colored in white in single channel and blue in the overlay images, translocates to mitochondria. **c** Mitochondrial and cytosolic fractions obtained from control and CCCP treated cells, as was described and analyzed in Fig. 2b, were immunoblotted for EndoB1 and OPTN, as indicated. **d** Hela OPTN/NDP52 knockdown cells were transfected with YFP-DAGR and mCherry-Parkin and treated with CCCP as indicated. EndoB1 localization was assessed by an EndoB1 antibody and was presented in pseudo color white as **b**. Note that translocation of EndoB1 to mitochondria was impaired in OPTN/NDP52 knockdown cells (bottom panel). **e** Cytosolic and mitochondrial fractions purified from control and OPTN/NDP52 knockdown cells treated with CCCP were immunoblotted for antibodies, as indicated. Note that mitochondrial EndoB1 levels were reduced in OPTN/NDP52 knockdown cells. **f,g** Hela cells were transfected with an EndoB1 siRNA, followed by transfection of expression plasmids for mCherry-Parkin and YFP-DAGR, and CCCP treatment at 10 µM for 9 hrs. Cells with mitochondrial YFP-DAGR were quantified from **f** (*P<0.05, Student’s t-test). Scale bar = 25 µM and zoom is 5x. Note that knockdown of EndoB1 suppressed mitochondrial DAG production.

## Discussion

Large-scale mitophagy activated by Parkin and CCCP treatment involves elaborate steps including tagging and transporting impaired mitochondria to the MTOC, followed by active dispersion to generate autophagible mitochondrial units, and de novo assembly of autophagosomes to restrict the dispersed mitochondria. In this report, we presented evidence that Parkin coordinates these complex events, at least in part, by stimulating local lipid remodeling on impaired mitochondria. The sequential production of mitochondrial PA and DAG activates actin-dependent dispersion of aggregated mitochondria and local autophagosome production, thereby enabling the efficient disposal of a large population of impaired mitochondria.

Our data show that Parkin orchestrates mitochondrial lipid remodeling via two enzymes, PLD2 and Lipin-1. Co-immunoprecipitation analysis indicates that Parkin can associate with PLD2 in response to mitophagy activation (Supplementary Fig. 2c). We, therefore, propose that the production of PA entails stable mitochondrial interaction of Parkin, which recruits PLD2 to the impaired mitochondria. The subsequent conversion of PA to DAG further requires Parkin ubiquitin ligase activity, indicating that Lipin-1, by itself, is not able to efficiently convert mitochondrial PA to DAG. Unlike PLD2, we did not detect apparent recruitment of Lipin-1 to Parkin-tagged mitochondria (data not shown). Instead, our data suggest that ubiquitin-binding autophagy receptors, OPTN and NDP52, deliver one or more co-factors to ubiquitinated mitochondria to enable Lipin-1-dependent DAG production. One such a co-factor we identified is EndoB1, which has an intrinsic PA binding activity ^41^. Our evidence suggests that EndoB1 is delivered by OPTN-positive Golgi-vesicles to ubiquitinated mitochondria (Fig. 6), where the mitochondrion-localized EndoB1 might bridge Lipin-1 and mitochondrial PA to produce DAG. Therefore, Parkin utilizes both ubiquitin-independent and -dependent process to orchestrate mitochondrial lipid remodeling.

The analysis of mitophagy phenotypes in Lipin-1-deficient cells has uncovered two inter-connected activities for mitochondrial DAG: activation of F-actin assembly and production of autophagosomes. It was previously observed that dispersion of perinuclear mitochondrial aggregates precedes autophagic sequestration of impaired mitochondria ^11^. Our data now provide evidence that the dispersion of mitochondrial aggregate is facilitated by the assembly of the actin cytoskeleton surrounding “aggregated” mitochondria (Fig. 3f, Middle Panels). Either inactivation of Lipin-1 or treatment with an actin polymerization inhibitor, cytochalasin B, effectively suppress mitochondrial de-aggregation (Fig. 3f, Bottom Panels and Fig. 3h). The actin cytoskeleton could, in principle, enable mechanical force production to separate individual mitochondria from the perinuclear cluster-a scenario similar to the actinomyosin-dependent de-aggregation of perinuclear protein inclusions, the aggresomes ^33^. Indeed, cortactin, a key co-factor for F-actin assembly, is required for the clearance of both aggresomes and mitochondrial aggregates ^12, 43^. Evidence also showed that active mitochondrial division mediated by Drp1 is necessary for efficient mitophagy^10^. Drp1-mediated mitochondrial fission also requires the cortactin-dependent actin cytoskeleton and involves EndoB1^44–46^. These findings support a model whereby mitochondrial DAG produced by EndoB1 and Lipin-1 nucleates local F-actin assembly to enable mitochondrial fission and dispersion.

Our analysis also revealed that Parkin-dependent DAG promotes mitophagosome production (Fig. 4). Accordingly, local production of DAG on Parkin- and ubiquitin-tagged mitochondria could, in principle, simultaneously activate focal mitochondrial fission/dispersion and de novo autophagosome production - providing a coupling mechanism to ensure “condemned” mitochondria are efficiently “restricted” by autophagosomes. Supporting the instructive role of DAG in autophagy activation, a cell-permeable DAG (DPG) can stimulate LC3-positive vesicle production in Lipin-1 KD cells (Fig. 4d-f). DPG treatment also markedly restored mitophagy in OPTN/NDP52 KD cells (Fig. 5c-d). The finding revealed that one key function of autophagic receptors in mitophagy is to stimulate DAG production. Interestingly, recent studies have revealed a surprising role of the actin cytoskeleton on autophagosome formation ^47, 48^. We propose that DAG-mediated F-actin assembly coordinates both the dispersion of mitochondria and local autophagosome assembly. Our data, however, do not exclude the potential effect of mitochondrial DAG on proteasome activity during mitophagy.

How mitochondrial DAG activates actin assembly remains to be determined. By mass spectrometry-based lipid analysis, we have detected a dramatic accumulation (>10 fold) of selected mitochondrial DAG species on mitochondria tagged for autophagy in response to CCCP treatment (Supplementary Fig. 6). This finding not only confirmed the production of mitochondrial DAG during mitophagy but also revealed that most prominently induced DAG species were those with long acyl chains (C38:4 and C38:3 as well as 36:2 and 36:1) while C32 and C34 species were not induced (Supplementary Fig. 6). Both C38:4 DAG (1-stearoyl-2-arachidonoyl-sn-glycerol; SAG) and C36:2 (1-stearoyl-2-linoleoyl-*sn*-glycerol; SLG) are potent PKC activators. These findings raise the possibility that mitochondrial DAG plays a signaling role and locally activates PKC on Parkin-tagged mitochondria to affect mitophagy. We further note that many LC3-positive autophagosomes produced during mitophagy are also labeled by the DAGR (Fig. 4b, Supplementary Fig. 3b), suggesting that DAG might have additional functions in the later stage of mitophagy. Future studies will be required to test these possibilities.

Perinuclear mitochondrial inclusions have been observed in both PD patients and a mouse PD model^22, 23, 49^. Perinuclear protein inclusion, the Lewy body, is a defining pathological feature of PD^50^. Despite their prevalence, the physiological purpose of concentrating “cellular junk” to the perinuclear region remains uncertain. Our findings suggest that perinuclear OPTN-associated and Golgi-derived vesicles are involved in mitochondrial DAG production and mitophagy. As the Golgi complex is normally located at the MTOC/perinuclear region, retrograde transport of impaired mitochondria and protein aggregates would place them close to the Golgi complex or Golgi-derived vesicles. Such an arrangement, in principle, could ensure a ready access of damaged mitochondria or protein aggregates to the OPTN/NDP52 vesicles for autophagic processing. Interestingly, the endoplasmic reticulum (ER) was proposed to play a critical role in mitophagy initiated at focally damaged mitochondria^20^. We surmise that when the production of damaged mitochondria exceeds the capacity of local autophagic degradation machinery (**peripheral autophagy**)-for example under chronic pathological conditions associated with neurodegenerative disease-the “overflowed” ubiquitin- and p62-tagged mitochondria would become engaged and transported by dynein motors to access the Golgi-associated vesicles and perinuclear autophagy machinery (**central autophagy**). Accordingly, the prevalence of perinuclear inclusions in PD patients could reflect the active role of the Golgi complex/vesicles in managing excessive cellular junk that accumulates in ailing neurons. Interestingly, many PD-associated PARK genes have been linked to Golgi integrity or Golgi-associated vesicular trafficking ^51–56^. It is tantalizing to speculate that Golgi dysfunction might be a risk factor for PD and that impaired central autophagy would weaken the clearance of protein aggregates or mitochondria, resulting in perinuclear inclusions.

In conclusion, our study has uncovered PLD2-EndoB1-Lipin 1 dependent mitochondrial lipid remodeling as a critical event in Parkin-dependent autophagic clearance of impaired mitochondria. Indeed, aberrant accumulation of mitochondria has been noted in Lipin 1 and EndoB1 knockout mice ^38, 57^, although the state of mitochondria in PLD2 knockout mice remains to be characterized. The effect of exogenous DPG to substitute for endogenous DAG in stimulating autophagosome production and mitophagy suggests that specific DAG-mimetics might improve mitochondrial QC in ailing neurons. Similar to impaired mitochondria, we have found that photo-damaged lysosomes targeted for autophagic clearance ^58^ also accumulated the PA reporter (W.Y.Y, Unpublished observation). An association of DAG on salmonella targeted for xenophagy has similarly been reported^59^. Thus, our study suggests that local lipid remodeling might be a conserved mechanism to implement autophagy-dependent organelle quality control (QC) and antibacterial defense.

## Materials and Methods

### Antibodies and Reagents

The following primary antibodies were used: anti-Tom20 (Santa Cruz sc-11415), anti-cytochrome c (BD Bioscience 556432), anti-GAPDH (Cell Signaling 14C10, 2118), anti-actin (Sigma AC-15, A1978), anti-EndoB1 (R&D, AF7456), anti-Tim23 (BD Bioscience), anti-Mfn2 (Santa Cruz sc-50331), anti-Mfn1 (Santa Cruz), anti-LC3 (MBL International), anti-FLAG (M2 Sigma F7425), anti-lipin-1 (Cell Signaling 5195), anti-PLD2 (Scbt sc-515744), Citrate synthetase (GTX110624, GeneTex), Hsp60 (Cell Signaling, 4870). Secondary antibodies were from Jackson Immunochem or Invitrogen.

The following reagents were used: DMSO (Sigma D8418), carbonyl cyanide 5-chlorophenylhydrazone (CCCP) (Sigma), PLD2 inhibitor VU 0364739 hydrochloride (Tocris 4171), Cytochalasin B (Sigma), and 1,2-dipalmitoyl-sn-glycerol (Sigma D9135)

### Cell Culture

Hela and HEK293T cells were obtained from Duke Cell Culture Facility (Durham, NC, USA). The cells were cultured in Dulbecco’s modified Eagle’s medium (DMEM; GIBCO-11995) supplemented with 10% fetal bovine serum and 1 × antibiotics (penicillin, 10,000 UI/ml and streptomycin, 10,000 UI/ml). These cell lines have been authenticated by STR DNA profiling and validated to be *mycoplasma*-free and before being frozen by the Duke Cell Culture Facility (Durham, NC, USA). All cells were maintained at 37`°`C and 5% CO_2_.

### Plasmids and Transfection

The following plasmids/siRNAs/shRNAs were used: GFP-Parkin and mutants as previously described^12^; mcherry-Parkin^20^; GFP-Raf1-PABD and CFP/YFP-DAGR^26^; mcherry-optineurin^14^; YFP-PLD1 and YFP-PLD2 (Gifts from W.G. Zhang); RFP-LifeAct (ibidi). mCherry-Raf1-PABD was generated by sub-cloning the PABD domain from GFP-Raf1-PABD to mCherry2(C1). Expression plasmids for WT and catalytic mutant mouse Lipin-1 were obtained from Addgene. Lipin-1 siRNA 5’-GAAUGGAAUGCCAGCUGAA-3’ and 3’-UUCAGCUGGCAUUCCAUUC-5’ (Invitrogen; HSS118307 (Sigma); optineurin siRNA (Invitrogen 4392420), NDP52 siRNA 5’-UUCAGUUGAAGCAGCUCUGUCUCCC-3’^60^, EndoB1 siRNA 5’-UGUUUAUACGACUUGGAGCUU-3’ and 3’-AAGCUCCAAGUCGUAUAAACA-5’ (Invitrogen), control siRNA (Ambion). Expression plasmids were transfected in HeLa and YFP-Parkin HeLa using Xtreme Gene 9 (Roche) according to manufacturer’s directions. Cells were transfected and treated 24-48 hours later. shRNA plasmids were transfected as stated above, but cells were not treated until 3-5 days later. siRNAs were transfected using RNAi MAX (Invitrogen) according to the manufacturer’s directions and cells treated 48-72 hours later.

### Immunofluorescence microscopy and quantification

Cells were seeded on coverslips, transfected, treated as indicated, and fixed in 4% PFA for 15 minutes. Coverslips were rinsed in PBS, permeabilized with 0.2% Triton-X 100 in PBS for 5 minutes, blocked in 10% BSA for 20 minutes in a humidified chamber, and incubated in primary antibodies diluted in 10% BSA overnight at 4C, followed by secondary antibodies in 10% BSA for 30 minutes. Coverslips were mounted on slides using Fluromount G (Southern Biotech). Slides were analyzed on a Leica SP5 confocal microscope using 100x/1.4-0.70NA or 40x oil objective (Leica Plan Apochromat). Z-stack images were acquired using the Leica LAS AF program software and maximum projections used in analyses and figures. Changes to brightness and contrast were performed in ImageJ. Number and size of vesicles were measured and quantified using Particle Analysis module in Image J.

For all immunofluorescent quantifications, at least 35 to 50 cells were counted for 2-3 separate experiments. Graphs represent means +/-SEM. For comparison of two conditions, a two-tailed, unequal student’s t-test was performed with p<0.05 considered significant. For all other comparisons, a one-way or two-way ANOVA was conducted.

### Preparation of mitochondrial fractions

The isolation of mitochondria from cultured cells was performed by following manufacturer’s protocol (#89874, ThermoFisher). In brief, after CCCP treatment, the cells harvested from a 10 cm plate (∼3 x 10^4^ cells) were first resuspended in reagent A. Cell membrane was then lysed using a reagent-based method by adding reagent B. After removing cell debris by low-speed centrifugation, the mitochondrial fractions were collected in pellets with high speed centrifugation. The remaining supernatant is cytosol fraction. Mitochondrial fractions were further subject to Western blots or lipidomic analysis.

### Western blots

HeLa or YFP-Parkin HeLa were seeded onto 10 cm or 6 cm dishes, transfected as outlined above, and treated as described. For western blot analysis, whole cell lysates were collected using 170mM NETN buffer, incubated at 4°C for 30 minutes, spun, supernatant collected, and protein concentration measured using the BCA assay (Thermo Scientific). Samples were normalized, diluted in XT buffer with reducing agent (BioRad), and boiled for 5 minutes at 100°C. 4-20% TGX gels (BioRad) were used, followed by transfer to nitrocellulose membranes. Primary antibodies were added to 2% BSA, incubated overnight at 4C, followed by rinsing, secondary antibody incubation for 1 hour at room temperature, rinsing, and developing using either ECL PicoWest (Thermo Scientific) or ECL Pro Lightning (Perkin-Elmer). Films were scanned, cropped, and adjusted for brightness and contrast in Photoshop. Density of bands was measured using Image J.

### Lipid extraction and LC/MS analysis

Lipid extraction of purified mitochondria was performed using a modified Bligh-Dyer method as previously decried ^61^. For LC/MS analysis, the dried lipid extracts were dissolved in chloroform/methanol (2:1, v/v) and injected for normal phase LC/MS analysis on an Agilent 1200 Quaternary LC system equipped with an Ascentis Silica HPLC column (5 μm, 25 cm x 2.1 mm, Sigma-Aldrich, St. Louis, MO). Mobile phase A consisted of chloroform/methanol/aqueous ammonium hydroxide (800:195:5, v/v/v); mobile phase B consisted of; mobile phase C consisted of. The elution program consisted of the following: 100% mobile phase A (chloroform/methanol/aqueous ammonium hydroxide; 800:195:5, v/v/v) was held isocratically for 2 min and then linearly increased to 100% mobile phase B (chloroform/methanol/water/aqueous ammonium hydroxide; 600:340:50:5, v/v/v/v) over 14 min and held at 100% B for 11 min. The LC gradient was then changed to 100% mobile phase C (chloroform/methanol/water/aqueous ammonium hydroxide; 450:450:95:5, v/v/v/v) over 3 min and held at 100% C for 3 min, and finally returned to 100% A over 0.5 min and held at 100% A for 5 min. The LC eluent (with a total flow rate of 300 μl/min) was introduced into the ESI source of a high resolution TripleTOF5600 mass spectrometer (Sciex, Framingham, MA). Instrumental settings for negative ion ESI/MS analysis of lipid species were as follows: IS= - 4500 V; CUR= 20 psi; GSI= 20 psi; DP= −55 V; and FP= −150 V. The MS/MS analysis used nitrogen as the collision gas. Data analysis was performed using Analyst TF1.5 software (Sciex, Framingham, MA).

### Data availability

All data supporting the findings of this study are available from the authors upon reasonable request. The source data underlying Figs 2d, 2f, 3b, 4e, 4f, 5b and 5d and Supplementary Figs 1a and 5d are provided as a Source Data file. Uncropped images of Western blots are shown (Supplementary Fig. 7).

## Acknowledgements

We thank Drs. M. Frohman and I. Kojima for PA and DAG reporters, W. G. Zhang for PLD1/2-YFP, E.F Holzbaur for OPTN-Cherry, and V. Bennett for TGN38-GFP expression plasmids. Dr. M Frohman for PLD6 KO MEFs. We thank Dr. T. Slotkin for the advice on the statistical analysis and YS Gao on the image analysis. This work was supported by 2R01-NS054022 (NIH) to T.-P.Y.

## Author contributions

K.L.S and T.P.Y. conceived the project initially. C.C.L, J.Y., M.D.K. and K.L.N designed and performed the majority of the experiments. Z.Q performed the lipidomic analysis. C.W.H. performed the live-cell imaging experiments. C.H.L. assisted with reagent preparation and experiments. T.P.Y. J.T.C and W.Y.Y supervised the work. N.V. and K.L.L. collaborated in the discussion and provided critical reagents. C.C.L., J.Y., M.D.K and T.P.Y. wrote the manuscript.

## Abbreviations List

OMM: Outer-mitochondrial membrane
IMS: inter-membrane space
IMM: inner-mitochondrial membrane
PA: phosphatidic acid
DAG: diacylglycerol
OPTN: optineurin
EndoB1: endophilin b1
CCCP: carbonyl chloro-m-phenylhydrazine
mitoPLD: phospholipase D 6
PINK1: PTEN-inducible kinase 1

