## Supplementary information for "Parkin coordinates mitochondrial lipid remodeling to execute mitophagy"

**Lin et al.**

### Lin_ Supplementary Figure 1

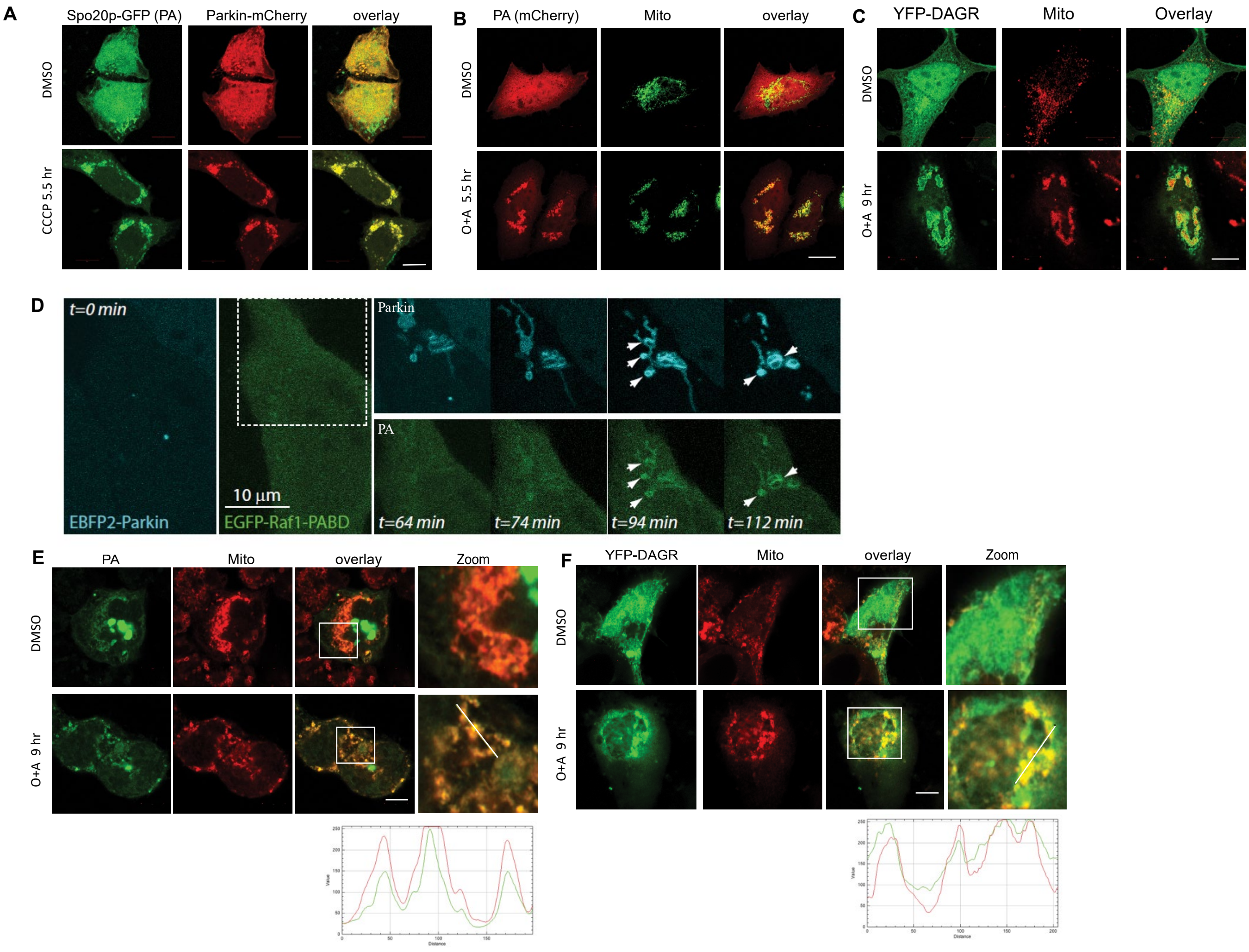


**Supplementary Figure 1. Mitophagy activation stimulates mitochondrial PA and DAG production. (A)** Hela cells were transfected with the Spo20p-GFP PA reporter and Parkin-mCherry followed by CCCP treatment for 5.5 hrs. Note that CCCP treatment led to the recruitment of Spo20p-GFP to Parkin-positive mitochondria. **(B-C).** HeLa cells were transfected with Raf1-PABD-mCherry PA reporter or YFP-DAGR with Parkin-FLAG followed by antimycin (4 mM) and oligomycin (10 mM; O+A) treatment to activate mitophagy. Note that Raf1-PABD-mCherry PA reporter (B) and YFP-DAGR (C) both translocated to mitochondria (TOM20) after the treatment. **(D)** A Hela cell expressing KR-dMito and EBFP2-parkin (cyan) were 559 nm illuminated to induce mitophagy. Images were acquired at indicated time points after illumination. Note that PA accumulation on Parkin-positive mitochondria induced by photodamage. (t=94 mins and t=112 mins). **(E-F)** SH-SY5Y cells were transfected with (E) the Raf1-PABD-GFP PA reporter or (F) YFP-DAGR followed by antimycin and oligomycin (10 mM) or DMSO treatment, as indicated. Both reporters became enriched at the mitochondria (TOM20). Line scan analysis (Image J software) under the images indicate colocalization between the lipid reporters (green) and mitochondria (red) corresponding to the lines drawn in the images. Scale bar = 10 µm.

### Lin_ Supplementary Figure 2

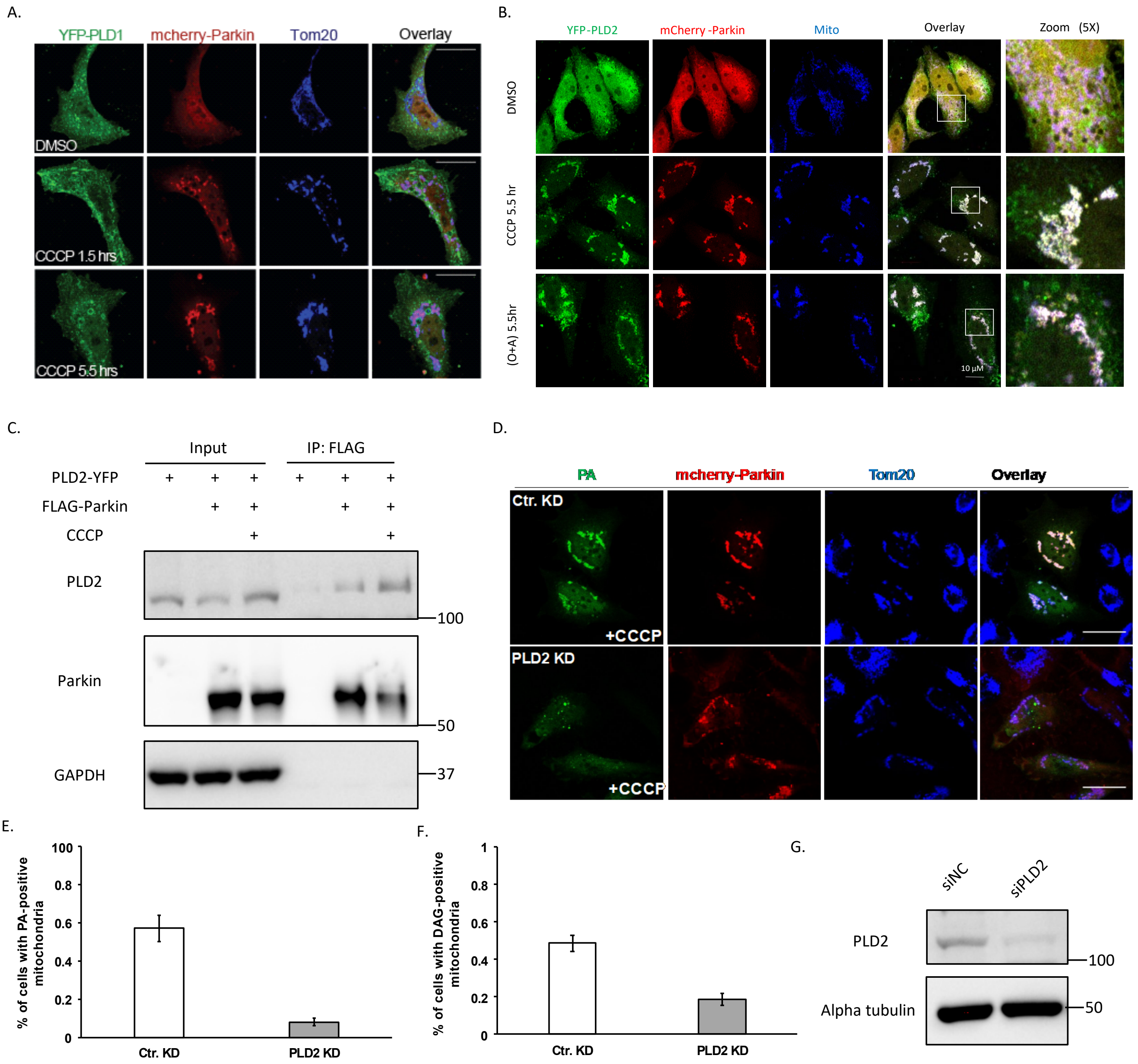

**Lin_ Supplementary Figure 2**

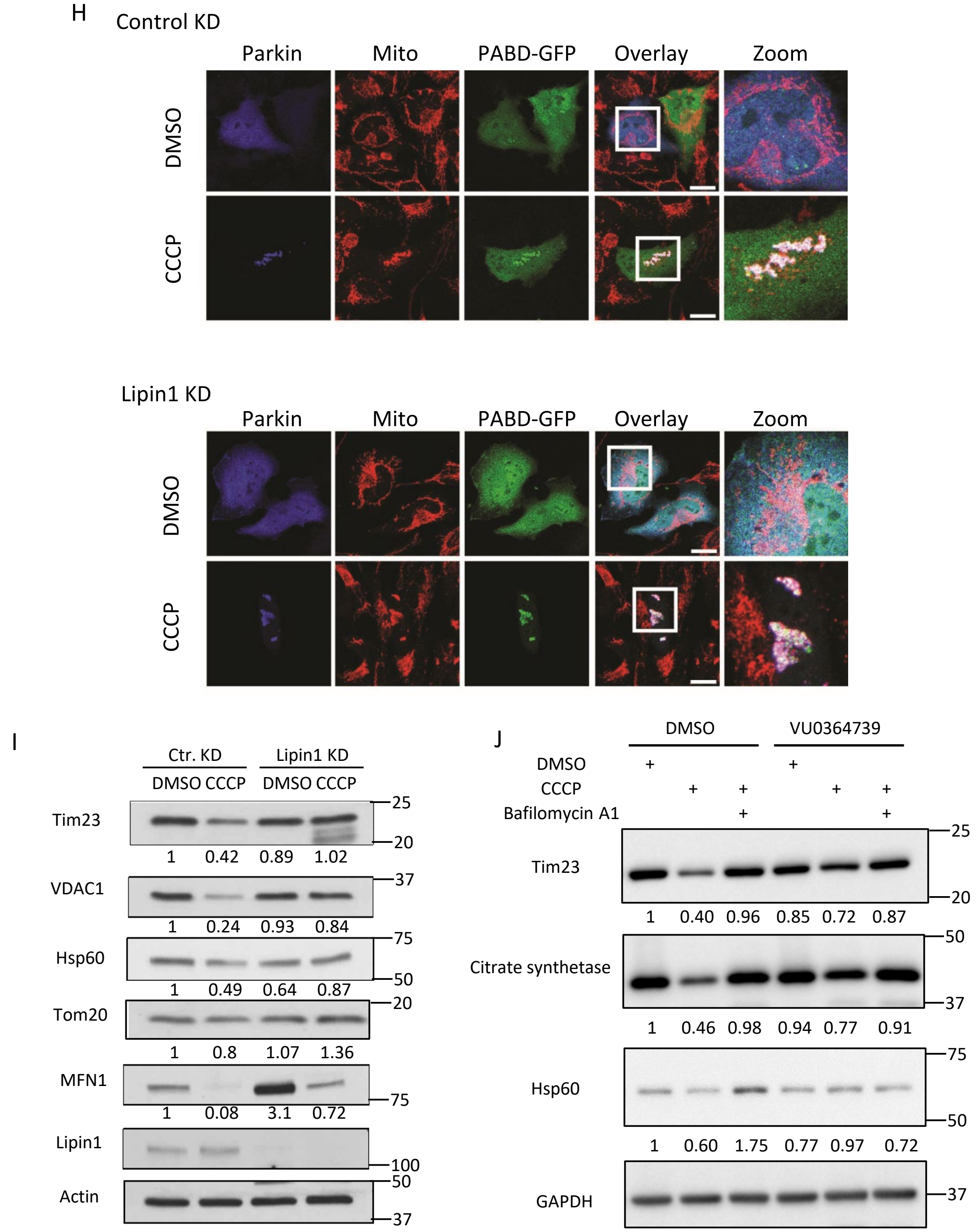


**Supplementary Figure 2. PLD1 is not translocated to mitochondria and Lipin-1 is dispensable for PA production.** (**A**) Hela cells were transfected with YFP-PLD1 and mcherry-Parkin followed by CCCP (10 µM) treatment at indicated time points. Note that PLD1 is not translocated to mitochondria upon CCCP treatment. (**B**) Hela cells were transfected with YFP-PLD2 and mCherry-Parkin followed by CCCP or oligomycin/antimycin treatment for indicated time. Note that PLD2 became colocalized with Parkin and mitochondria (TOM20) under both treatment conditions. **(C)** HEK-293T cells expressing PLD2-YFP and FLAG-Parkin cDNA were treated with CCCP for 5.5 hours. Parkin was then pulled down by a FLAG antibody from cell lysates and blotted with a PLD2 antibody. Note that PLD2 interaction with Parkin was enhanced by CCCP. **(D-G)** Hela cells were transfected with siRNA for PLD2 followed by expression plasmids for mCherry-Parkin and the PA or GFP reporter, and subject to DMSO or CCCP treatment, as indicated. PLD2 knockdown inhibited both mitochondrial PA reporter (D-E) and DAG reporter accumulation (F). PLD2 knockdown efficiency was confirmed by immunoblotting (G). **(H)** Hela cells were transfected with Lipin-1 siRNA followed by plasmids for FLAG-Parkin and the PA reporter and treated with DMSO or CCCP, as indicated. Note that PA accumulated on mitochondria (TOM20) in Lipin-1 knockdown cells. Scale bar is 25 µM and zoom is 3x. **(I)** Hela cells stably expressing Parkin-YFP were treated with Lipin1 siRNA and subjected to CCCP (10 µM) for 18 hrs. **(J)**. Hela cells stably expressing Parkin-mCherry were treated with PLD2 inhibitor (VU0364739, 3µM) and subjected to CCCP (10 µM) and Bafilomycin A1 (1µM, lysosomal inhibitor) for 18 hrs. Note that PLD2 inhibition rescued mitochondrial protein degradation. The band intensity of mitochondrial proteins relative to control untreated conditions were determined by Image J.

### Lin_ Supplementary Figure 3

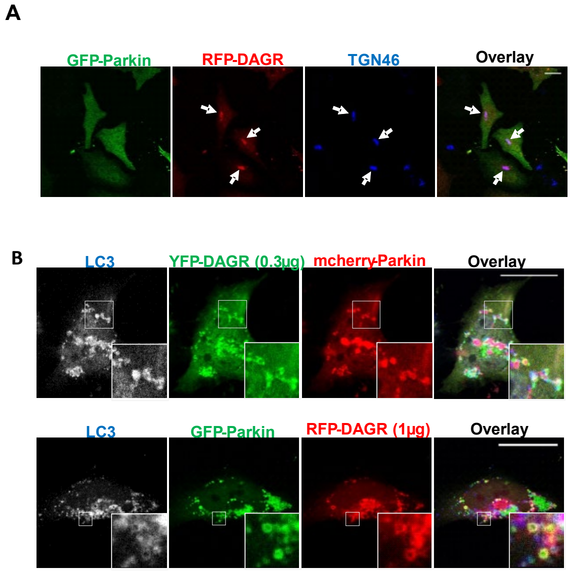


**Supplementary Figure 3. RFP-DAGR and YFP-DAGR sequesters similar mitophagosome structure and the effect of Lipin-1 knockdown on autophagosome production. (A)** Hela cells were transfected with GFP-Parkin and RFP-DAGR. Trans-golgi was assessed by immunostaining with a TGN46 antibody. Note that RFP-DAGR is colocalized with TGN46. Scale bar is 25 μM and zoom is 5x (A-top panel and B), 10x(A-bottom panel). **(B)** Hela cells were transfected with either GFP-Parkin and 1 μg of RFP-DAGR (bottom panels of A) or mcherry-Parkin and 0.3 μg of YFP-DAGR (top panels of A) followed by CCCP treatment. Autophagosomes were assessed by immunostainining of LC3 antibody. Note that YFP-DAGR formed the similar LC3-positive structure sequestering dispersed mitochondria as RFP-DAGR did in Fig. 4 (B top panel).

### Lin_ Supplementary Figure 4

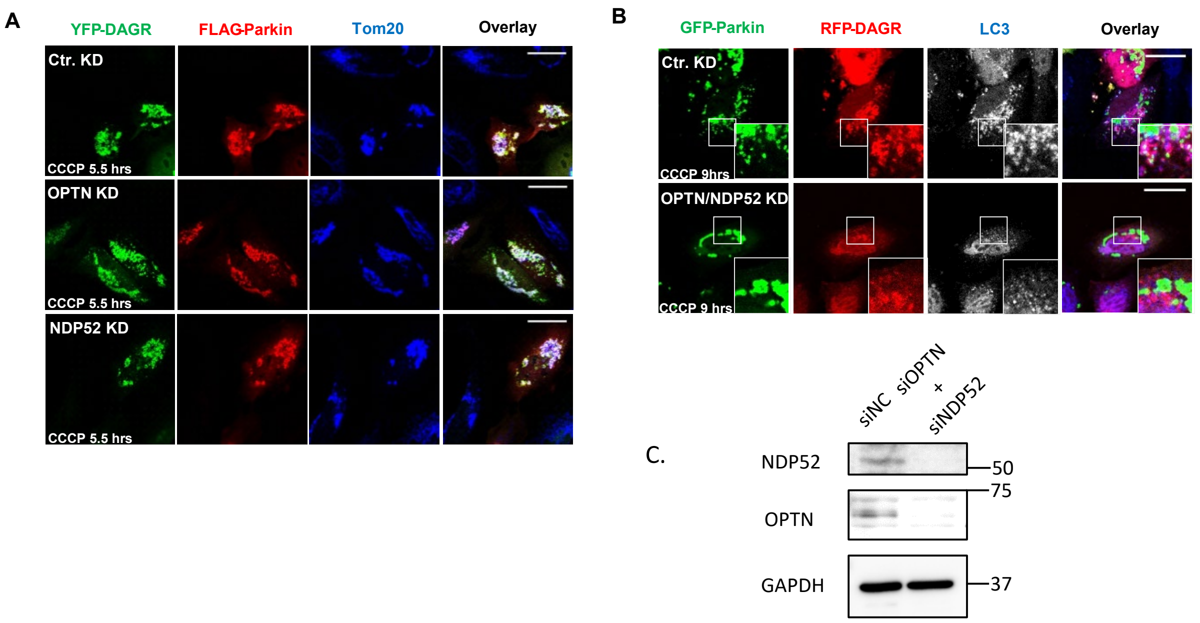


**Supplementary Figure 4. The effect of optineurin (OPTN) or NDP52 single knockdown on mitochondrial DAG production and double knockdown on mito-autophgosome formation. (A)** Hela cells were transfected with control siRNA, OPTN siRNA or NDP52 siRNA. Knockdown cells were transfected with the DAGR-YFP reporter and FLAG-Parkin followed by CCCP treatment. Note that DAGR accumulated on mitochondria (Tom20) in both OPTN knockdown cells and NDP52 knockdown cells. **(B)** OPTN and NDP52 double knockdown cells were transfected with GFP-Parkin and RFP-DAGR and treated with CCCP at 10 μM for 9 hrs. Autophagosomes were assessed by immunostaining with a LC3 antibody. Note that DAG-positive LC3 vesicles production was reduced in OPTN and NDP52 double knockdown cells. Scale bar = 25 μM and zoom is 5x. **(C)** Knockdown efficiency for OPTN and NDP52 was confirmed by immunoblotting.

#### Lin_ Supplementary Figure 5

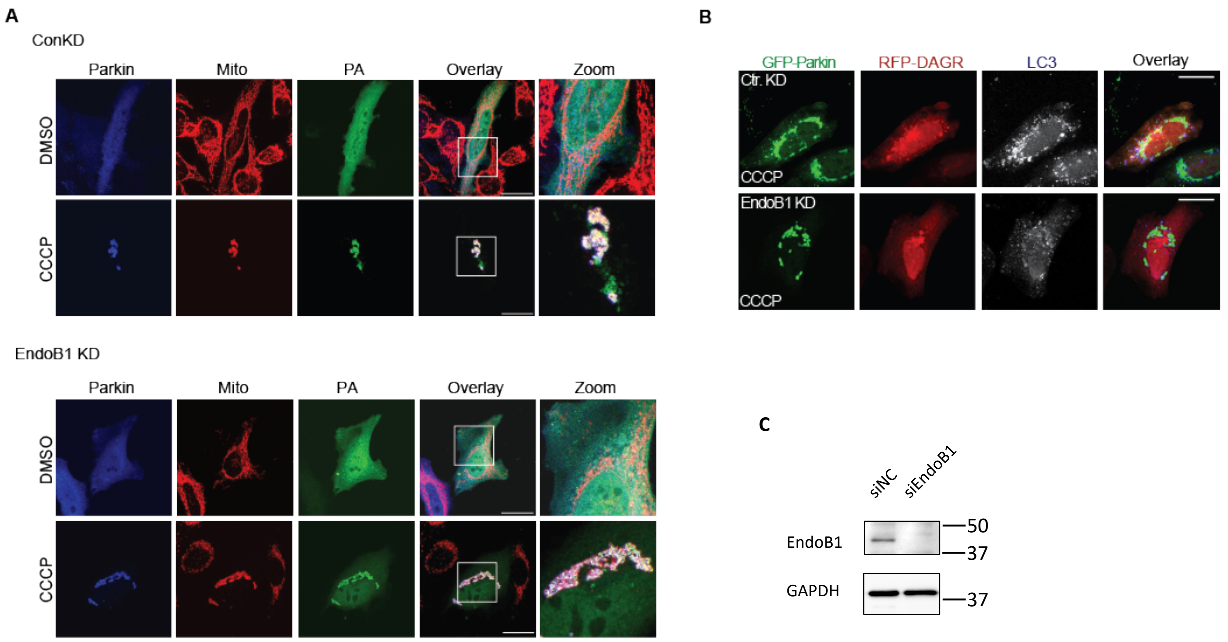


**Supplementary Figure 5. Effect of EndoB1 knockdown on mitochondrial PA and mito-autophagosome production.** (**A**) EndoB1 knockdown cells were transfected with either FLAG-Parkin and the PA reporter followed by CCCP treatment. Mitochondria was assessed by immunostaining of Tom20 antibody. Note that PA accumulated on mitochondria in EndoB1 knockdown cells. (**B**) EndoB1 knockdown cells were transfected with GFP-Parkin and DAG reporter RFP-DAGR followed by CCCP treatment. Autophagosomes were assessed by immunostaining of LC3 antibody. Note that DAG-positive LC3 vesicle production was mitigated in EndoB1 knockdown cells. Scale bar = 25 μM and zoom is 3x. (**C**) Control and EndoB1 KD cells were subjected to immunoblotting by EndoB1 and GAPDH antibodies, as indicated, to confirm knockdown efficiency.

#### Lin_ Supplementary Figure 6

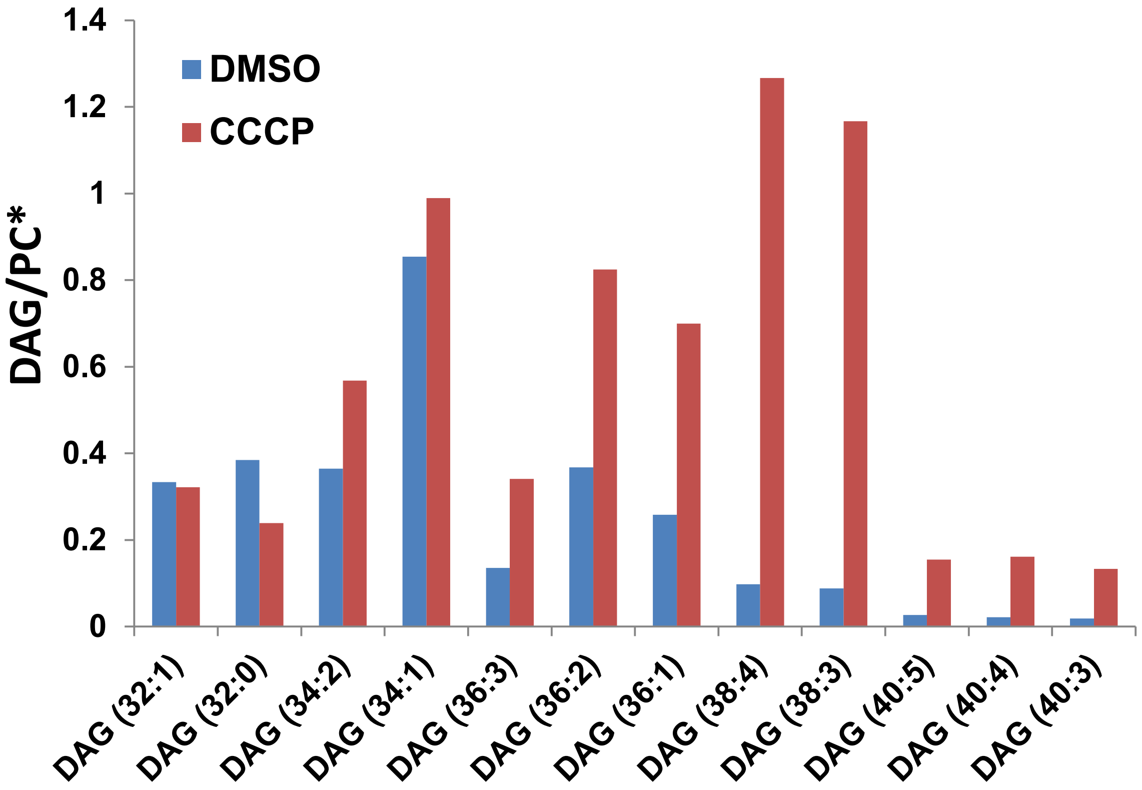


**Supplementary Figure 6. Selected DAG species are accumulated on mitochondria targeted for autophagy.** Mitochondria purified from control, and CCCP treated (5 hrs) Parkin expressing HeLa cells were subjected to the lipidomic analysis by LC/MS (see Material and Method for details). The relative abundance of DAG species was normalized by phosphatidylcholine (PC), which was not affected by CCCP. Note the dramatic accumulation of selected DAG species on mitochondria in response to CCCP.

### Supplementary Figure 7 Uncropped images of Western blot

**Fig 2b**

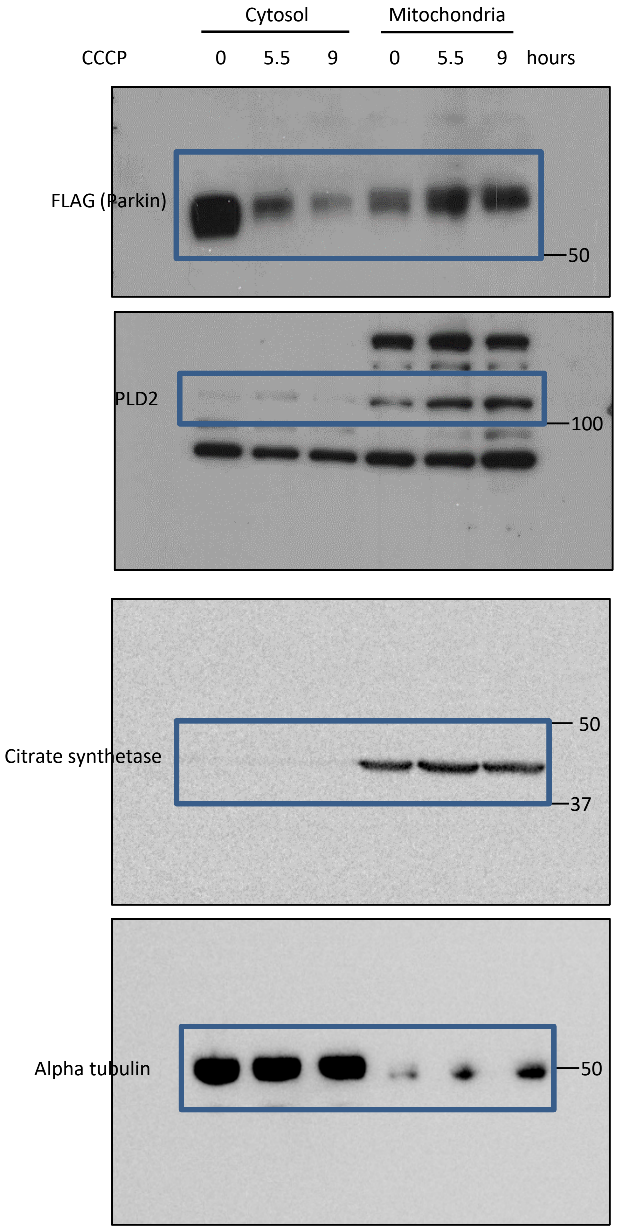

**Fig 3c**

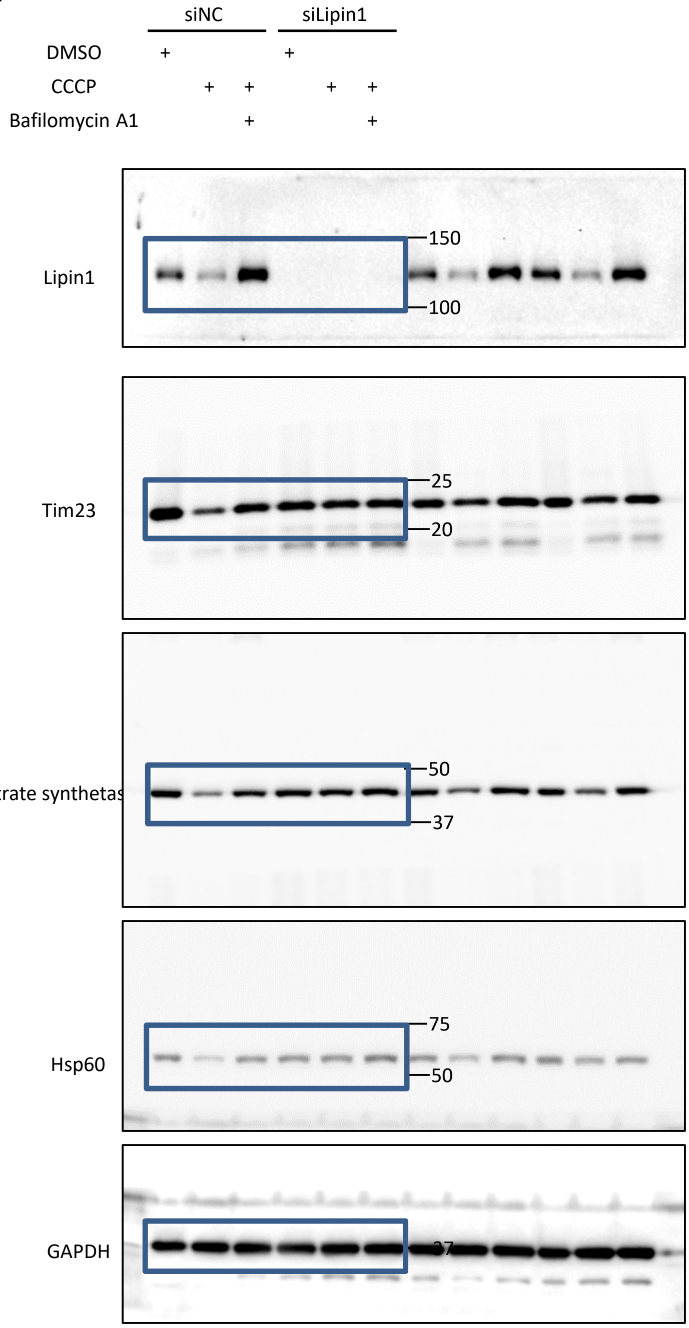
siNC
siLipin1
DMSO
CCCP
Bafilomycin A1
+
+
+
+
+
+
+
+
Lipin1
150
100
Tim23
25
20
Citrate synthetas
50
37
Hsp60
75
50
GAPDH
37

**Fig 6c**

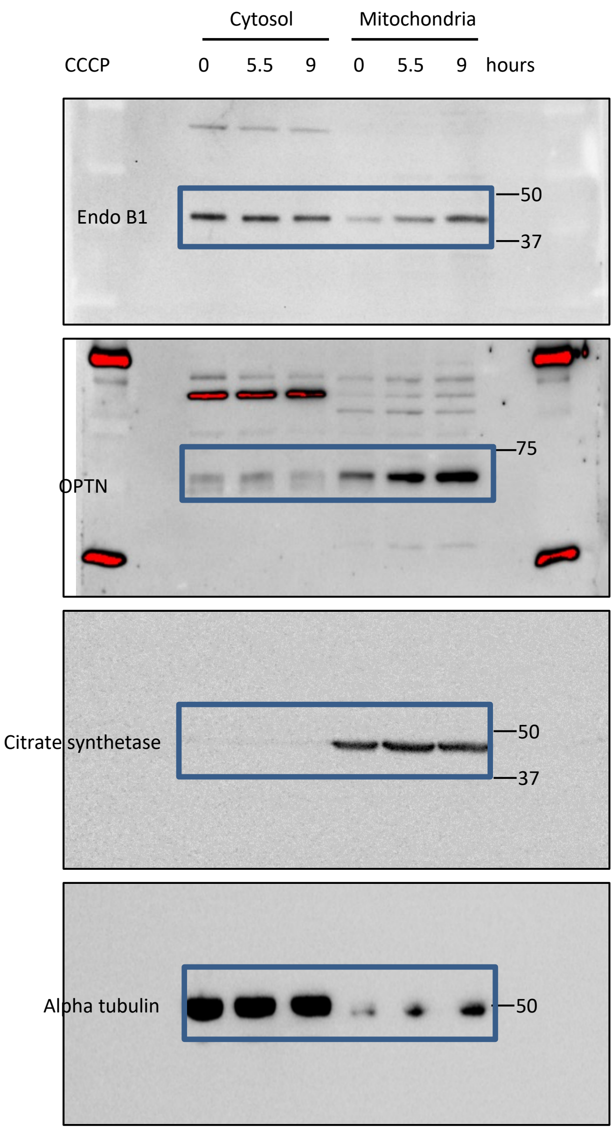
Cytosol
Mitochondria
CCCP
0
5.5
9
0
5.5
9
hours
Endo B1
50
37
OPTN
75
Citrate synthetase
50
37
Alpha tubulin
50

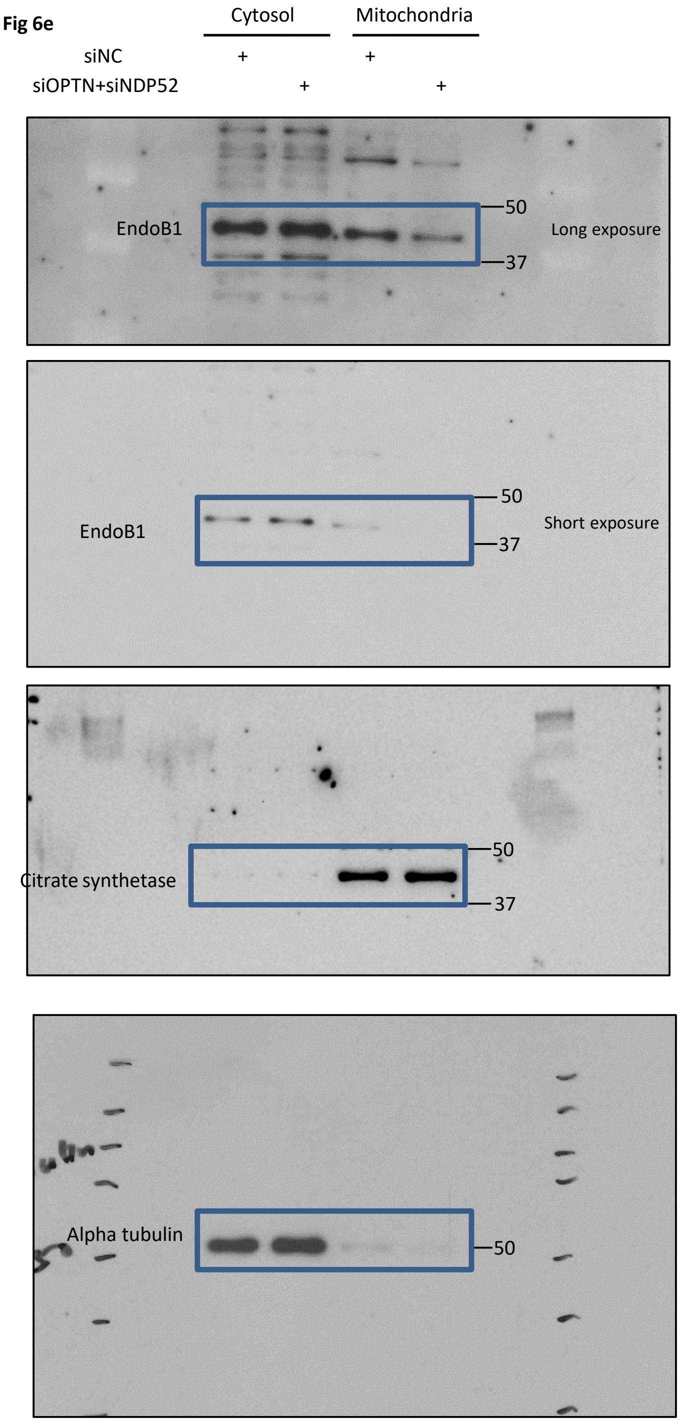
Fig 6e
Cytosol
Mitochondria
siNC
+
+
siOPTN+siNDP52
+
+
EndoB1
50
37
Long exposure
EndoB1
50
37
Short exposure
Citrate synthetase
50
37
Alpha tubulin
50

**Fig S2c.**

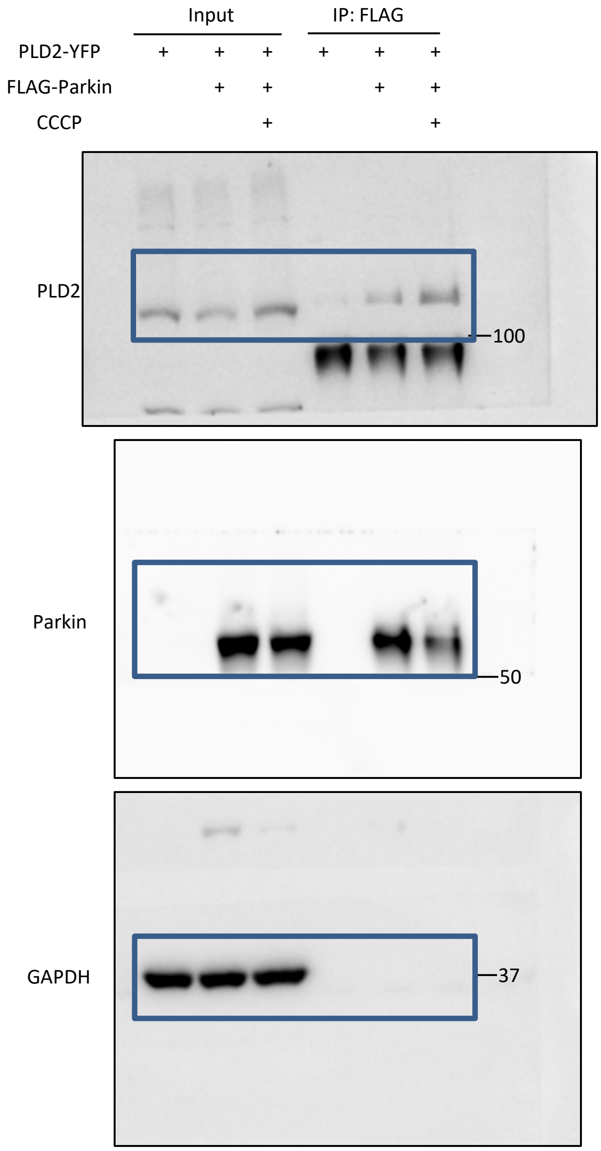


Fig S2g

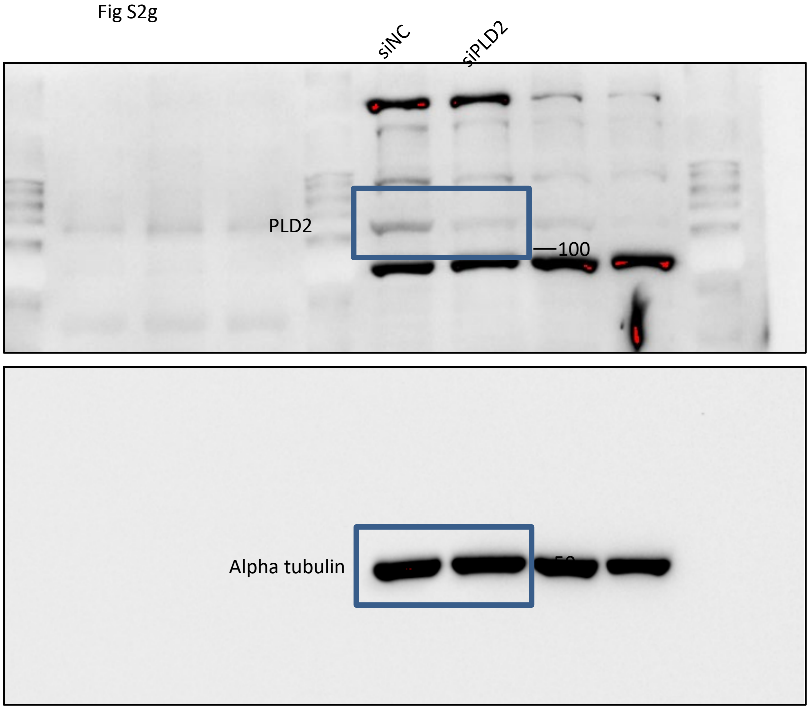

Fig. S2j

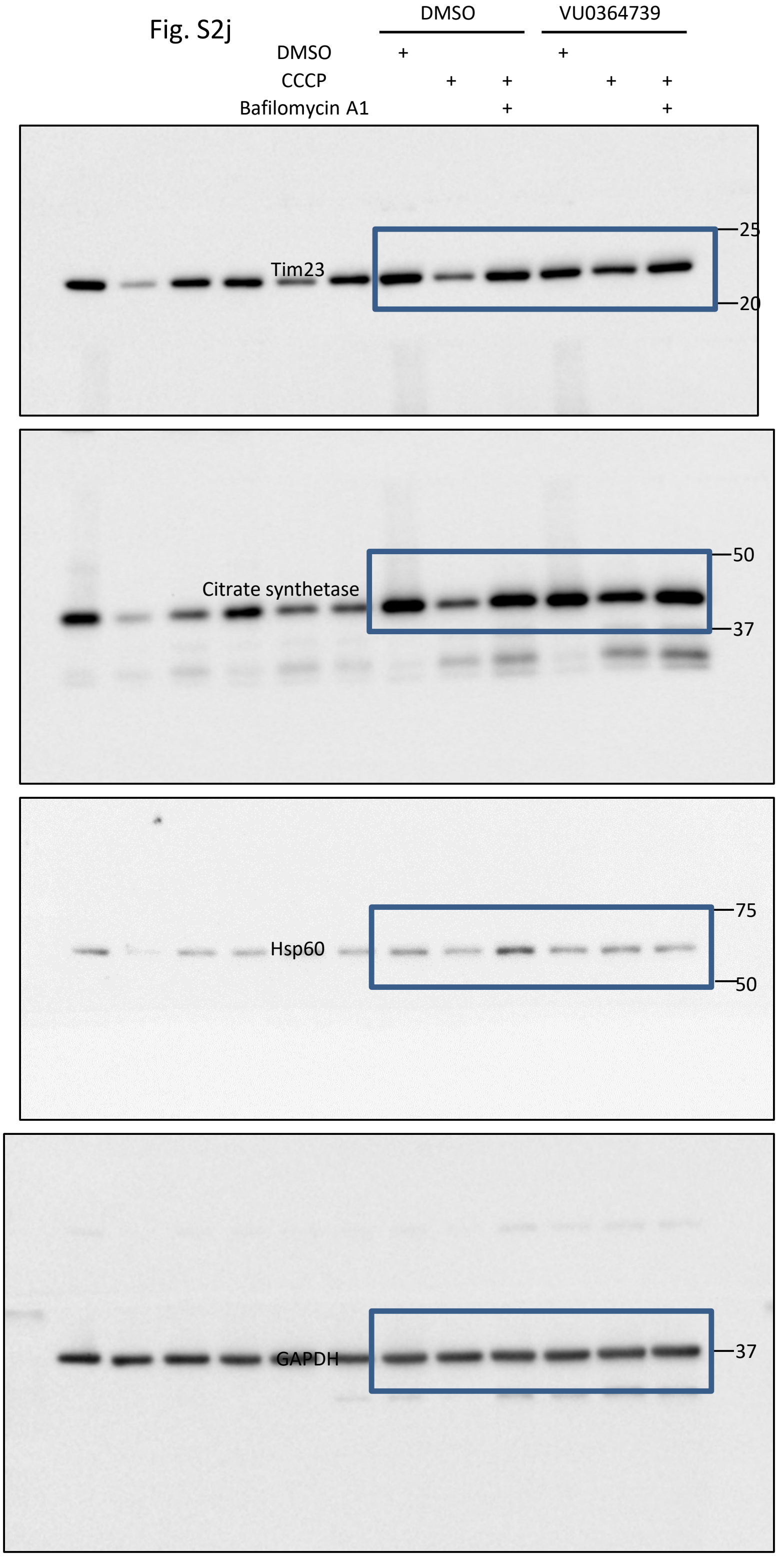
DMSO
VU0364739
DMSO
CCCP
Bafilomycin A1
Tim23
25
20
Citrate synthetase
50
37
Hsp60
75
50
GAPDH
37

Fig S4c

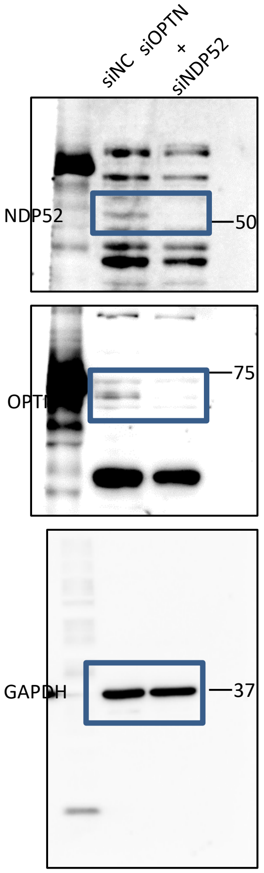


Fig S5c

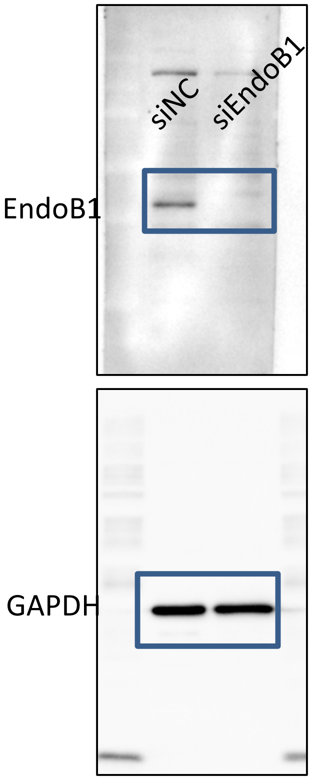
